# Full-length de novo viral quasispecies assembly through variation graph construction

**DOI:** 10.1101/287177

**Authors:** Jasmijn A. Baaijens, Bastiaan Van der Roest, Johannes Köster, Leen Stougie, Alexander Schönhuth

## Abstract

Viruses populate their hosts as a viral quasispecies: a collection of genetically related mutant strains. Viral quasispecies assembly is the reconstruction of strain-specific haplotypes from read data, and predicting their relative abundances within the mix of strains is an important step for various treatment-related reasons. Reference-genome-independent (“de novo”) approaches have yielded benefits over reference-guided approaches, because reference-induced biases can become overwhelming when dealing with divergent strains. While being very accurate, extant de novo methods only yield rather short contigs. The remaining challenge is to reconstruct full-length haplotypes together with their abundances from such contigs. We present Virus-VG as a de novo approach to viral haplotype reconstruction from pre-assembled contigs. Our method constructs a variation graph from the short input contigs without making use of a reference genome. Then, to obtain paths through the variation graph that reflect the original haplotypes, we solve a minimization problem that yields a selection of maximal-length paths that is optimal in terms of being compatible with the read coverages computed for the nodes of the variation graph. We output the resulting selection of maximal length paths as the haplotypes, together with their abundances. Benchmarking experiments on challenging simulated and real data sets show significant improvements in assembly contiguity compared to the input contigs, while preserving low error rates compared to the state-of-the-art viral quasispecies assemblers. Virus-VG is freely available at https://bitbucket.org/jbaaijens/virus-vg.

## 1 Introduction

The ensemble of genetically related, but different mutant viral strains that populate infected people are commonly referred to as *viral quasispecies* (Domingo *et al*., 2012). Each of these strains comes with its own genomic sequence (henceforth referred to as *haplotype*). The final goal of primary viral quasispecies analysis is the reconstruction of the individual haplotypes— optimally at full length—and also to provide estimates of their abundances. The unknown number of different, strain-specific haplotypes and their variance in abundance establish the theoretical issues that characterize viral quasispecies assembly. They explain why this form of assembly is difficult, despite the shortness of virus genomes. These issues are further accentuated by the fact that neither next-generation nor third-generation sequencing reads, by their combinations of error rates and length, allow for immediate reconstruction and abundance estimation of haplotypes (Beerenwinkel *et al*., 2012; Rose *et al*., 2016).

State-of-the-art approaches currently allow for two options: (**i**) full-length reconstruction of haplotypes based on statistical, usually *reference genome dependent* measures, or (**ii**) *de novo* reconstruction of (optimally haplotype-specific) contigs.

Approaches of type (**i**) assume that the sequencing reads are aligned to a reference genome and make use of model-based clustering algorithms (Zagordi *et al*., 2011; Barik *et al*., 2017; Ahn and Vikalo, 2018), Dirichlet process mixture models (Prabhakaran *et al*., 2014), hidden Markov models (Töpfer *et al*., 2013), sampling schemes (Prosperi and Salemi, 2012), or combinatorial methods (Knyazev *et al*., 2018), respectively. However, as was demonstrated in Baaijens *et al*. (2017); Töpfer *et al*. (2014), resorting to external auxiliary means (such as reference genomes) can bias the reconstruction procedure significantly.

Approaches of type (**ii**) comprise generic (meta)genome assemblers as well as specialized viral quasispecies assemblers, both of which are not helped by external measures (“*de novo*”) hence are not affected by external biases. Metagenome assemblers are designed to reconstruct multiple genomes simultaneously, but in viral quasispecies tend to collapse strains (Rose *et al*., 2016). It was further shown by Baaijens *et al*. (2017) that among generic de novo assemblers SPAdes (Bankevich *et al*., 2012) was the only approach to identify strain-specific sequences, however only in case of sufficiently abundant strains. *De novo* viral quasispecies assemblers (e.g. Hunt *et al*. (2015); Yang *et al*. (2012)) generally aim at constructing suitable consensus reference genomes, which may serve as a template for more finegrained studies (for example if curated reference genomes have become too divergent, which is a frequent scenario). To only methods that truly aim at *de novo* genome assembly *at strain resolution* are SAVAGE (Baaijens *et al*., 2017), MLEHaplo (Malhotra *et al*., 2016) and PEHaplo (Chen *et al*., 2018). However, the contigs produced by these methods, while strain-specific, in general do not represent full-length haplotypes.

We present Virus-VG, an algorithm that turns strain-specific contigs into full-length, strain-specific haplotypes, thus completing the de novo viral quasispecies assembly task. For that, we construct a variation graph from the contigs, without the help of a (curated) reference genome, where we use the contigs produced by SAVAGE (Baaijens *et al*., 2017). We obtain full-length haplotypes as a selection of maximal-length paths in the variation graph, each of which reflects a concatenation of subpaths associated with the input contigs. The selected paths are optimal in terms of differences between their estimated abundances and the read coverages computed for the nodes they traverse. Although our approach to quasispecies assembly using candidate path enumeration followed by abundance estimation is similar to Astrovskaya *et al*. (2011); Skums *et al*. (2013), these methods make use of read graphs rather than variation graphs and define optimality for path abundance estimation in a very different way.

Variation graphs are mathematical objects that have recently become very popular as reference systems for (evolutionarily coherent) collections of genomes (Paten *et al*., 2017). Using such genome structures instead of standard linear reference genomes has been shown to reduce reference bias (Dilthey *et al*., 2015; Paten *et al*., 2017) and to allow for efficient subhaplotype match queries (Novak *et al*., 2017) and haplotype modelling (Rosen *et al*., 2017). Methods presented so far for constructing variation graphs, however, have been focused on a linear reference genome as a point of departure. Here, we point out how to construct variation graphs *de novo*, by first sorting the contigs in an appropriate way and then making use of progressive multiple alignment techniques (`vg msga`, part of the `vg toolkit` by Garrison *et al*. (2017)). In this, we present an approach for full-length, high-quality reconstruction of the haplotypes of a viral quasispecies that is *entirely de novo*.

Our method depends on the enumeration of maximal-length paths in a variation graph, whose number is in the worst case exponential in the number of nodes of the graph. However, since all these paths enumerated are to respect the subpaths associated with the input contigs, their number will decrease on increasing contig length. Thanks to advances in sequencing technology, input contig length will inevitably increase, which points out that our method, as per its design, will be able to deal with future technological developments smoothly.

Benchmarking experiments demonstrate that Virus-VG yields substantial improvements over the input contigs assembled with SAVAGE in terms of spanning the full length of the haplotypes. Thereby, the increase in length comes at negligible or even no losses in terms of sequential accuracy compared to the input contigs. Further, we find our strain abundance estimates to be highly accurate. Finally, we find our method to (substantially) outperform alternative approaches, all of which are reference based—we recall that there are no alternative de novo approaches so far—both when working with bootstrap (i.e. assembled from the data itself) and curated reference genomes.

*Note on Related Work: RNA Transcript Assembly.* Many RNA transcript assemblers work in a similar way to Virus-VG: first enumerating all possible transcripts, then selecting an ‘optimal’ set of transcripts using various optimization methods (Feng *et al*., 2010; Li *et al*., 2011; Mezlini *et al*., 2013). This has led to variations of minimum path cover optimization problems that are—regarding a few relevant aspects—similar in spirit to the optimization problem we formulate (Bernard *et al*., 2014; Pertea *et al*., 2015; Rizzi *et al*., 2014; Tomescu *et al*., 2013; Trapnell *et al*., 2010). Most importantly, Rizzi *et al*. (2014) introduce node and edge abundance errors and Tomescu *et al*. (2013) show a minimum path cover with subpath constraints to be polynomially solvable. However, to the best of our knowledge, no method simultaneously employs both subpath constraints *and* abundance error minimization in its problem formulation. Moreover, applying these RNA transcript assemblers to the viral quasispecies problem is not so straightforward: a collection of reference genomes representing all possible haplotypes is required as input, while in our setting such information is not available.

## 2 Methods

**Notation.** A *variation graph* (*V, E, P*) is a directed graph that is constructed from a set of input sequences, which represent members of a (evolutionarily coherent) population of sequences. Each node *v* ∈ *V* is assigned to a subsequence seq(*v*). An edge (*u, v*) ∈ *E* indicates that the concatenation seq(*u*)seq(*v*) is part of one of the input sequences. *P* is a set of paths (a sequence of nodes linked by edges) that represent genome-specific sequences; thereby, *P* can, but need not, represent the input sequences themselves. A node *v* ∈ *V* with no incoming edges is called *source*. A node *v* ∈ *V* with no outgoing edges is called *sink*.

**Workflow.** Our method consists of two basic steps:

1. The computation of a ***contig variation graph*** *VG*′ = (*V*′, *E*′, *P*′) where each path *p* ∈ *P*′ represents an input contig. We refer to the path representing contig *c* as *p*(*c*). Together with *VG*′, we compute a function *a*′: *V*′ → ℝ where *a*′(*v*′) for *v*′ ∈ *V*′ represents the abundance of an individual node, measured by the amount of original reads (from which the contigs were computed) that align to seq(*v*′).
2. The transformation of *VG*′ into a ***genome variation graph*** *VG* = (*V, E, P*) where each path *p* ∈ *P* reflects a full-length haplotype. We also compute a function *a*: *P* → ℝ where *a*(*p*) for *p* ∈ *P* reflects the abundance of the haplotype represented by *p*. The set of paths *P* together with their abundances *a*(*p*) establish the final output of our method.

The input for determining *VG*′ in (**1**) are the contigs. For computation of *a*′, we make use of the original reads from which the input contigs were computed; one can determine the abundance *a*′(*v*′) of single node *v*′ ∈ *V*′ as the (length normalized) count of reads whose alignments touch upon *v*′.

The input for computation of *VG* and *a* in (**2**) are *VG*′ and *a*′. Since *V* ⊆ *V*′ and *E* ⊆ *E*′, as will become clear later, we can apply *a*′ also to nodes in *VG*. The computation of *VG* is established as the solution of an optimization problem that aims to determine full-length paths (paths formed by a concatenation of contigs of maximal length) such that the difference of path abundances *a*(*p*) and node abundances *a*′(*v*) for paths *p* of which *v* makes part is minimal. We emphasize here that the numbers *a*′(*v*) can be directly computed from the input, whereas the *a*(*p*)’s correspond to decision variables in an optimization problem.

We will describe the construction of the contig variation graph (**1**) in full detail in Section 2.1. The transformation into the (final) genome variation graph (**2**) is divided into two steps: (**a**) the *enumeration of candidate paths*, which is described in Section 2.2.1, and (**b**) the *solution of an optimization problem* that aims at selecting a subset of candidate paths through their path abundance values which are optimal in terms of being compatible with node abundances in Section 2.2.2. The complete workflow is illustrated in Figure 1.

**Figure 1:**
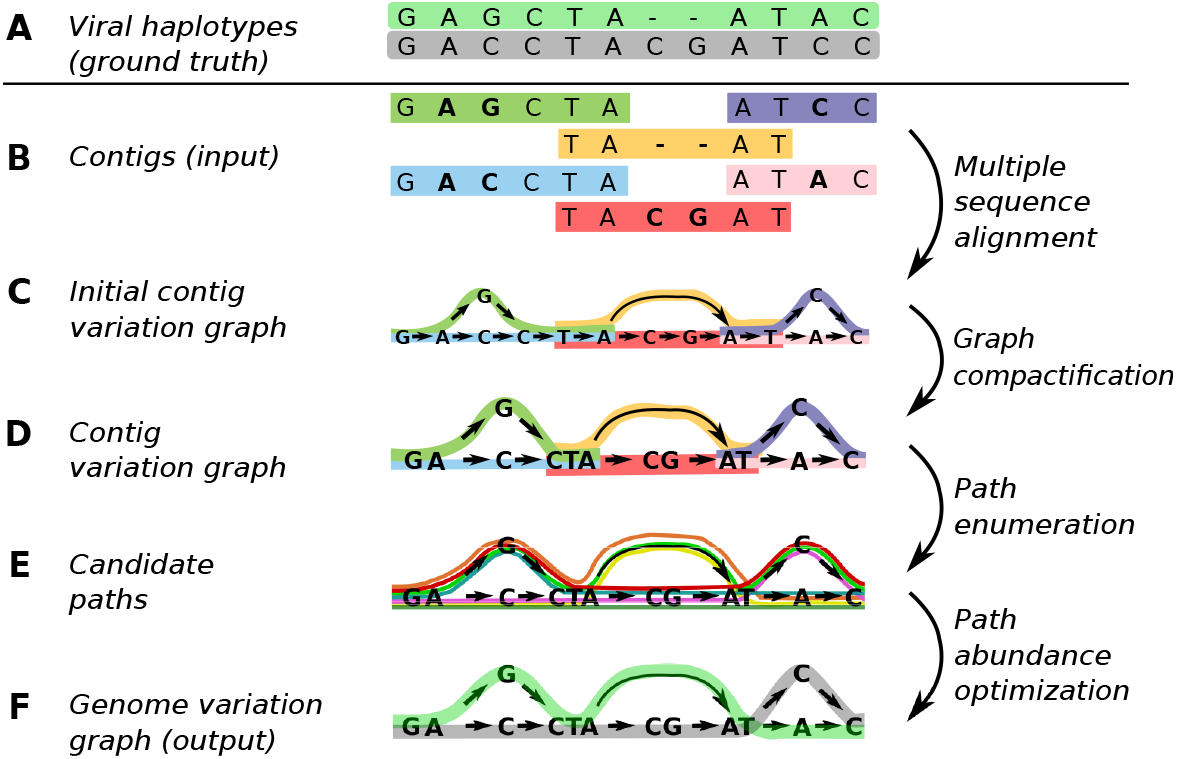
Virus-VG workflow. (A) Ground truth haplotypes from which the sequencing reads originate; (B) Input contigs, obtained by de novo assembly; (C) Initial contig variation graph created from input contigs using multiple sequence alignment; (D) Contig variation graph obtained by collapsing non-branching paths (compactification); (E) Candidate paths representing possible haplotypes; (F) Genome variation graph representing the haplotypes selected through path abundance optimization, capturing the viral quasispecies.

### 2.1 Contig variation graph construction

**Input**. The input is a data set of next-generation sequencing reads and a set of contigs assembled from them, for which we use the specialized de novo viral quasispecies tool SAVAGE (Baaijens *et al*., 2017). We assume that there are no contigs which are an exact subsequence of another contig, which applies for SAVAGE (and commonly applies for the output of many assembly programs); any such contigs are removed. The contig variation graph with its node abundances is constructed in three steps.

**Step 1: Multiple Sequence Alignment (MSA).** We construct the initial contig variation graph by building an MSA of the contigs using `vg msga` (Garrison *et al*., 2017), which progressively combines long sequences into a variation graph. For this construction to work on a collection of contigs that do not all cover the same region, the order in which the contigs are aligned and added to the graph is important: we need to add contigs such that they overlap the existing graph as much as possible, to avoid creating more disjoint components than necessary. Here, we sort the contigs by starting with the longest contig, then iteratively selecting the contig with longest possible overlap with any of the previously selected contigs, until all contigs have been selected. For finding all pairwise overlaps between contigs we use `minimap2` (Li, 2018). Determining the best sorting heuristic in terms of speed and quality is subject to future work. After sorting the contigs, we apply `vg msga`; the resulting MSA is represented as a variation graph and for every contig the corresponding path through the graph is stored.

**Step 2: Compactification and contig-path construction.** We compactify the initial contig variation graph in a similar fashion as in the construction of a compacted de Bruijn graph (Mäkinen *et al*., 2015). The absence of branches on a path ensures that every source-sink path has to traverse it at full length. Therefore, each non-branching path (*v*_*i*_1__,…,*v_i_k__*) can be merged (contracted) into a single node 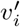, with in-neighbors 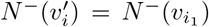 and out-neighbors 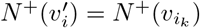. Also the contributing contig sets of *v*_*i*_1__,…,*v_i_k__* are taken together and stored in the new node 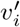. Note that after this step, a node can represent a sequence instead of a single nucleotide.

In addition, we determine for each contig c the sub-path *p*(*c*) in this (compacted) graph that represents *c*. Let *p*(*c*) = (*v*_*i*_1__,…,*v_i_k__*) be this sub-path. Note that due to the compression step, the sequence seq(*c*) represented by a contig *c* might only be a subsequence of its path sequence seq(*v*_*i*_1__)…seq(*v_i_k__*). However, this does not bear any consequence on the definition of any haplotype the contigs make part of.

The resulting compacted graph, together with the contig paths *P*′ is our contig variation graph *VG*′ = (*V*′, *E*′, *P*′), illustrated in Figure 1, panel D.

**Step 3: Node abundance.** We finally compute *a*′: *V*′ → ℝ, which assigns *node abundances a*′(*v*′) to nodes *v*′ ∈ *V*′ of the contig variation graph. These node abundances *a*′(*v*′) reflect the *average base coverage* of the piece of sequence seq(*v*′). For computation of *a*′(*v*′) we make further use of the `vg-toolkit` (Garrison *et al*., 2017), which allows to align the original sequencing reads to our contig variation graph. The abundance *a*′(*v*′) is calculated as the sum of all bases in all reads that align with seq(*v*′), divided by the length of seq(*v*′).

### 2.2 From contig to genome variation graph

The input for the following procedure is the contig variation graph *VG*′ = (*V*′, *E*′, *P*′) together with *a*′: *V*′ → ℝ that we have just described. The procedure for constructing the *genome variation graph VG* = (*V, E, P*) from *VG*′ and *a*′ consists of three steps. *First*, we compute a set of *candidate paths*, which are all maximal length paths in (*V*′, *E*′) that are “concatenations” of paths from *P*′. *Second*, we select a subset of candidate paths that are optimal with respect to a *minimization problem*, which provides us with the final, maximal-length paths *P* and path abundances *a*: *P* → ℝ. *Third*, we remove nodes and edges from (*V*′, *E*′) that are not traversed by paths from *P*, which yields the final graph (*V, E*). Since only paths in *P* are supposed to reflect true haplotypes, any node not being included in a haplotype is most likely a sequencing artifact or an assembly error (or it belongs to a missing strain). The third step is a straightforward procedure. We will describe the first two steps in more detail in the following.

#### 2.2.1 Candidate path generation

The goal is to compute the set of all paths through (*V*′, *E*′) that are maximal-length *concatenations* of paths from *P*′, where we understand a concatenation of two paths as the merging of them along a common substring. Thereby, this common substring is a suffix of the first path and a prefix of the second path. We will refer to these paths as *candidate paths P*_cand_ in the following (see also Figure 1, Panel E). Generating candidate paths happens in five steps outlined below.

**Step 1: Trimming paths** *p* ∈ *P*′. Due to common issues in contig computation, uncorrected sequencing errors are often located on the extremities of the contig. We therefore shorten all paths *p* ∈ *P*′ by their extremities and remove the tails if these contain nodes *v*′ for which *a*′(*v*′) is below a given threshold. By default, we allow to trim paths *p* ∈ *P*′ by a removal of nodes that together amount to no more than *τ* = 10bp on either end.

**Step 2: Enumerating pairwise concatenations.** We allow concatenating pairs of paths with matching suffix-prefix pairs. In more detail, let *p*_1_, *p*_2_ ∈ *P*′, represented by series of nodes (*u*_1_,…,*u_m_*) and (*v*_1_,…,*v_n_*) from *V*′. Then *p*_1_ can be concatenated with *p*_2_, written *p*_1_ →_*c*_ *p*_2_, if for some *l* we have *u*_*m*−*l*+1_ = *v*_1_, *u*_*m*−*l*+2_ = *v*_2_,…,*u_m_* = *v_l_*, that is, the suffix of length *l* of *p*_1_ matches the prefix of length *l* of *p*_2_.

In order to enable correction of persisting sequencing errors, we further consider to concatenate pairs of paths *p*_1_, *p*_2_ which do have one or more non-matching nodes, but only under the following condition. Let *u*\*:= *u*_*m*−*l*+*i*_ ≠ *v_i_* =: *v*\* be the respective non-matching nodes in *p*_1_, *p*_2_ respectively, then only if min{*a*′(*u*\*), *a*′(*v*\*)} < *α*, where *α* is a user-defined threshold, we concatenate *p*_1_ and *p*_2_. This threshold reflects the minimal node abundance *a*′(*v*) for which we trust node *v*; for more details, see Appendix A.

**Step 3: Removing concatenations lacking physical evidence.** Subsequently, we remove concatenations *p*_1_ →_*c*_ *p*_2_ if there are *q*_1_, *q*_2_ such that *q*_1_ →_*c*_ *q*_2_, *q*_1_ →_*c*_ *p*_2_, *q*_2_ →_*c*_ *p*_2_, but there is no *q*_3_ for which *p*_1_ →_*c*_ *q*_3_ and *q*_3_ →_*c*_ *p*_2_ and there is *q*_4_ such that *p*_1_ →_*c*_ *q*_4_. The situation reflects that the concatenation of paths *q*_1_ →_*c*_ *p*_2_ enjoys corroborating physical evidence, provided by *q*_2_, while there is no such corroborating evidence for the concatenation *p*_1_ →_*c*_ *p*_2_. At the same time, *p*_1_ concatenates well with *q*_4_ such that the removal of *p*_1_ →_*c*_ *p*_2_ does not turn *p*_1_ into a dead end.

**Step 4: Enumerating maximal length paths *P*_cand_.** In this step, the pairwise concatenations from step 2 that remain after step 3 are combined to paths of maximal length. This is achieved through a breadth-first search type procedure. We maintain a set of *active paths P*_act_, which is the set of paths to be extended in the current iteration. We also maintain a set of *maximal paths P*_max_ that reflects the set of maximal length paths collected.

1. **Initialization:** We determine all *p* ∈ *P*′ for which there are no *q* →_*c*_ *p* and put them both into *P*_act_ and *P*_max_.
2. **Iteration:** We replace each *p* ∈ *P*_act_ with all *q* ∈ *P*′, for which *p* →_*c*_ *q* without *q*\* such that *p* →_*c*_ *q*\* →_*c*_ *q*. Simultaneously, we extend each 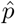 ∈ *P*_max_ that ends in *p*, by appending *q* (while respecting the overlap). In case *q* is already part of 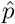, we do not append *q* to 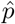 but instead add *q* as a new path to *P*_max_, thereby breaking any possible loops due to repetitive elements.
3. **Output:** If for all *p* ∈ *P*_act_ there are no *q* with *p* →_*c*_ *q*, we output *P*_max_ as our candidate path set *P*_cand_. The enumeration algorithm lists all candidate paths in time linear in the output size. This can be (in the worst case) exponential in the number of paths *p* ∈ *P*′, depending on the structure of the contig variation graph.

**Step 5: Correcting paths for errors.** After enumerating all candidate paths, we apply a final correction step to every such path. Since we allow concatenating paths from *P*′ where suffix-prefix pairs do not match in all nodes (see Step 2), we may have positions in candidate paths where contig paths *p* ∈ *P*′ do not agree on the underlying sequence. All such ambiguous positions refer (by construction) to low abundance nodes *v*′ (i.e., *a*′(*v*′) < *α*). We resolve the ambiguity by selecting the node *v*\* from all contributing paths *p* ∈ *P*′ with maximal abundance *a*′(*v*\*).

#### 2.2.2 Minimization for haplotype selection and abundance estimation

**Input.** For this final part of the method, the input is the set of candidate haplotype paths *P*_cand_ and the node abundances *a*′(*v*). In general this set of paths is much larger than the actual number of haplotypes, so *P*_cand_ will contain many false haplotypes. Here we filter them out by estimating the abundance *a*(*p*) for each path (haplotype) *p* ∈ *P*_cand_ through solving a minimization problem. In a subsequent step, haplotype paths with an abundance below a user defined threshold will be removed as being most likely false haplotypes. This leaves the set of haplotypes to be output.

**Determining path abundances** *a*(*p*). We determine path abundance values *a*(*p*) for every *p* ∈ *P*_cand_, such as to minimize the sum of or, equivalently, the average of node abundance errors. Let *f*(*x, y*) be an error function to be chosen later. Then for node *v* the node abundance error is defined as the value of *f*(*x, y*) with *x* the node abundance *a*′(*v*) and *y* the sum of the abundances of the haplotype paths going through the node *v*, which is 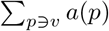. Recall that the node abundance values *a*′(*v*) are obtained from read alignments to the contig variation graph (Section 2.1, Step 3). The objective then becomes minimizing the sum of the node abundance errors over all nodes *v* ∈ *V*′:

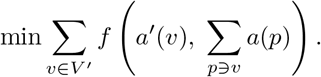

We need to add non-negativity constraints *a*(*p*) ≥ 0 on the path abundances. Since we have already taken all subpath constraints into account when enumerating the candidate haplotype paths, the minimization problem does not need any further constraints.

Note that the effectiveness of this objective function depends heavily on the error function used as well as the correctness of node abundances *a*′(*v*). These abundance values are not exact measurements, but based on read alignments to the graph as described above; coverage fluctuations can thus lead to under- or overestimated node abundance values. In this case, a simple linear objective function is preferred over a quadratic error function, because the former allows big errors in certain nodes to be compensated by small errors in other nodes. We also observed that normalizing the errors w.r.t. the true node abundance does not improve results, because this means that errors in nodes with low abundance values are penalized very strongly. For this reason, we use the error function *f*(*x, y*) = |*x* − *y*| in our objective and the optimization problem becomes

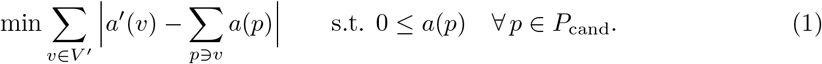

This is a convex programming formulation, which can, by a standard transformation, easily be linearized and solved using an LP solver.

**Output: haplotype selection and final abundances.** The outcome of the minimization problem (1), yields for each *p* ∈ *P*_cand_ an optimal abundance value *a*\*(*p*). We now select the set of haplotype paths as output of the procedure, by removing any haplotypes with an estimated abundance below a user defined threshold *γ*. In other words, as output we give the set *P* = {*p* ∈ *P*_cand_ | *a*\*(*p*) ≥ *γ*} (Figure 1, panel F). After this haplotype selection step, we redo the optimization step on the selected haplotype paths (prefixing *a*(*p*) to 0 for every path *p* with *a*\*(*p*) < *γ*), thus ensuring that our final abundance estimates are as accurate as possible.

**Note on related work.** The minimization problem we are treating here can be considered as a combination of problems presented in Rizzi *et al*. (2014) and Tomescu *et al*. (2013). The combination of these problems would require an unambiguous way to have subpath abundances contribute to cumulative abundances on the nodes. It is not immediately evident how to do so. In our setting it is straightforward how path abundances *a*(*p*) contribute to the estimated abundances of the nodes on the paths. Exploring these relationships is interesting future work.

## 3 Results

We present results for Virus-VG on four challenging simulated data sets and one real benchmark. We compare our method with the viral quasispecies assemblers ShoRAH (Zagordi *et al*., 2011) and PredictHaplo (Prabhakaran *et al*., 2014), which are widely approved and state-of-the-art in terms of full-length reconstruction of viral haplotypes. On a shorter region (HIV pol gene) we also compare our method to aBayesQR (Ahn and Vikalo, 2018) and PEHaplo (Chen *et al*., 2018). Although a comparison to the RNA transcript assemblers from Rizzi *et al*. (2014) and Tomescu *et al*. (2013) would be interesting, this is not so straightforward: these methods require as input a collection of reference genomes representing all possible transcripts (or in our case, viral haplotypes). Since we do not have such information, we could not apply these methods to our data.

For shell commands, parameters to be set, their default choices, and further reasoning, see the Supplementary Material.

### 3.1 Data sets

For evaluating correctness of our algorithm and benchmarking experiments on full-length viral genomes, we selected the two most challenging simulated data sets (HCV, ZIKV) presented by Baaijens *et al*. (2017) and generated one additional data set (Poliovirus). These data sets represent typical viral quasispecies ultra-deep sequencing data and consist of 2×250bp Illumina MiSeq reads which were simulated using SimSeq^1^. In addition, we simulated mixtures of HIV strains, considering only the 3kb pol region, at various sequencing depths. Finally, we also consider a real Illumina MiSeq data set commonly used for benchmarking, referred to as the *labmix*. More details about all data sets are presented in the Supplementary Material.

*10-strain HCV mixture.* This is a mixture of 10 strains of Hepatitis C Virus (HCV), subtype 1a, with a total sequencing depth of approximately 20,000x (i.e. 400,000 reads). The haplotypes were obtained from true HCV genomes in the NCBI nucleotide database and have a pairwise divergence varying from 6% to 9%. Paired-end reads were simulated at relative frequencies between 5% and 13% per haplotype, i.e., a sequencing depth of 1000x to 4600x per haplotype.

*15-strain ZIKV mixture.* This is a mixture of 15 strains of Zika Virus (ZIKV), consisting of 3 master strains extracted from the NCBI nucleotide database and 4 mutants per master strain. The pairwise divergence varies between 1% and 12% and the reads were simulated at relative frequencies varying from 2% to 13.3%. The total sequencing depth for this data set is again 20,000x.

*6-strain Poliovirus mixture.* This is a mixture of 6 strains of Poliovirus (type 2), with a total sequencing depth of approximately 20,000x. The haplotypes were obtained from true Poliovirus genomes from the NCBI database. Paired-end reads were simulated at exponentially increasing relative frequencies of 1.6% to 50.8%.

*7-strain HIV mixture (pol gene).* This is a mixture of 7 HIV-1 pol gene sequences with a pairwise divergence of 1%, obtained by introducing random mutations into sequence D86068.1 from the NCBI database. Paired-end reads (2×250bp) were simulated for each of the 7 strains at relative frequencies between 0.5% and 61.5%. We created 3 data sets with sequencing depths of 500x, 1000x, and 5000x, respectively.

*Labmix.* This is a real Illumina MiSeq (2×250 bp) data set with an average coverage of 20,000x, sequenced from a mixture of five known HIV-1 strains (HXB2, NL4-3, 89.6, YU2, JRCSF) with relative strain frequencies between 10% and 30%. This data set was presented as a benchmark by Di Giallonardo *et al*. (2014) and is publicly available^2^. Currently, predictions of all methods, including our own, are hampered by highly repetitive regions such as the long terminal repeats on the HIV genome; see also Baaijens *et al*. (2017). Hence, we decided to follow Di Giallonardo *et al*. (2014) in removing these a priori by excluding any reads that map to these known repeat sequences.

### 3.2 Assembly evaluation criteria

We use QUAST (Gurevich *et al*., 2013) for evaluating our experiments and report the number of contigs, the fraction of the target genomes that was reconstructed, the N50 and NGA50 measures, and observed error rates. Here, the target genome consists of all true haplotypes known to be present in a sample; a base is considered reconstructed if there is at least one contig with an alignment to this base. The N50 measure, defined as the length for which the collection of all contigs of that length or longer covers at least half the assembly, gives an indication of assembly contiguity. The NGA50 measure is computed in a similar fashion, but only aligned blocks are considered (obtained by breaking contigs at misassembly events and removing all unaligned bases). This measure reports the length for which the total size of all aligned blocks of this length or longer equals at least 50% of the total length of the true haplotypes; the NGA50 value is undefined if a target coverage of 50% cannot be reached. Finally, the error rates we present are computed as the sum of the N-rate (i.e. ambiguous bases) and mismatch- and indel rates (compared to the ground truth), normalized by the number of assembled bases. Further details are presented in the Supplementary Material.

### 3.3 Improvements of final haplotypes over input contigs

The first two rows of Table 1a, SAVAGE and Virus-VG, display the statistics for the input contigs and the final, maximal-length haplotypes computed here, respectively, for the HCV data set. While SAVAGE presents 26 fragmented contigs, Virus-VG presents 10 full-length haplotypes, each of which represents one of the original haplotypes, thereby encompassing the 10 original haplotypes that established the basis for simulating reads. Further, Virus-VG covers 99.3% of the target genomes, similar to the original 99.4% provided by the input contigs, and these full-length haplotypes come at a negligible error rate of 0.001%. In summary, our approach yields near-perfect results on this (supposed to be challenging) data set.

**Table 1:**
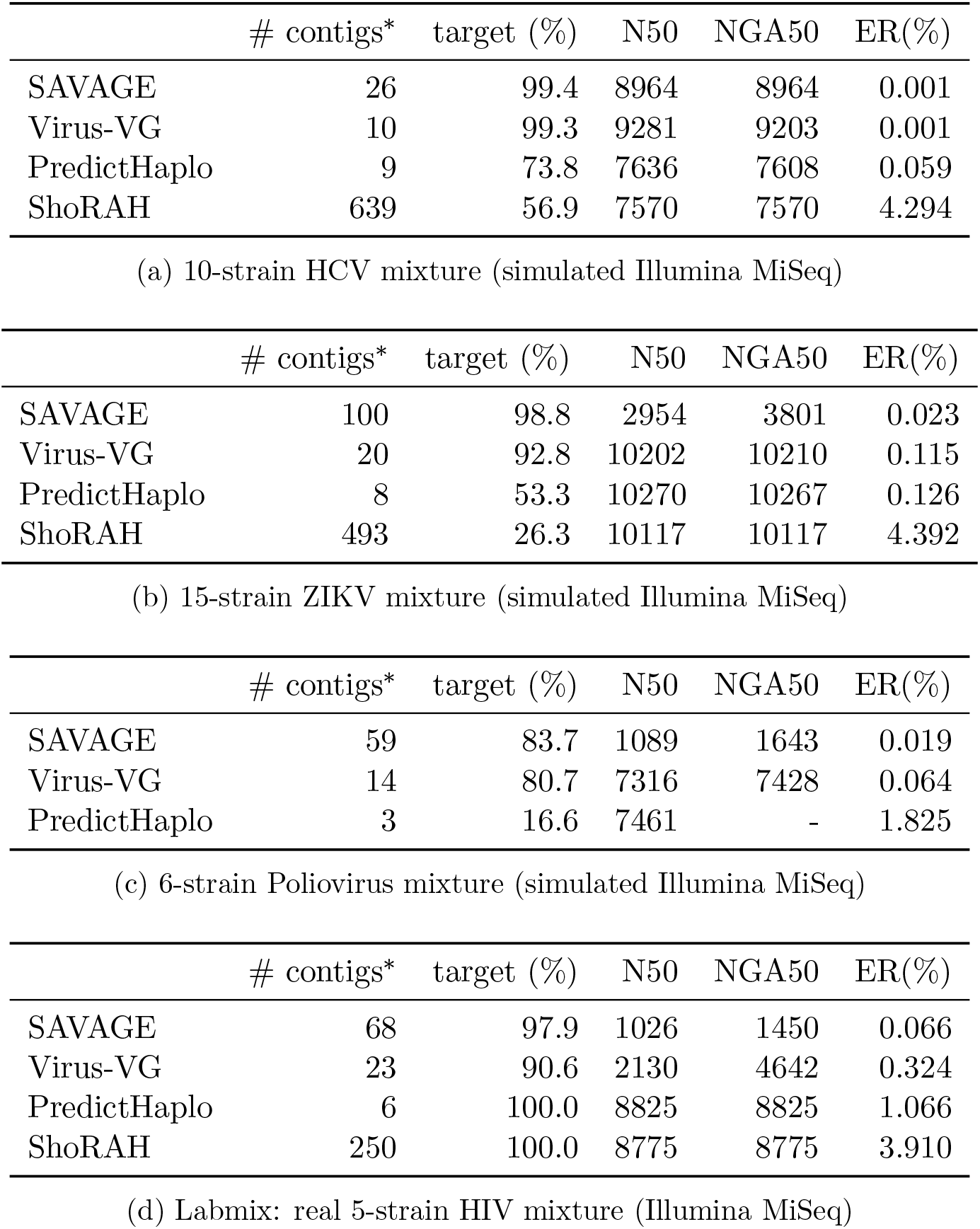
Assembly results per dataset. ER = Error Rate, computed as the sum of the fraction of ‘N’s (ambiguous bases) and the mismatch- and indel rates. ShoRAH could not process the Poliovirus data. *If contigs are full-length, this number reflects the estimated number of strains in the quasispecies.

For the 15-strain ZIKV data set (Table 1b) we again achieve substantial improvements in terms of haplotype assembly contiguity. We obtain 20 full-length haplotypes covering 14 out of 15 strains, while the original input contigs consisted of 89 highly fragmented, and relatively short sequences. As a result, we observe an NGA50 value of 10210 for Virus-VG, reflecting full-length haplotypes, compared to an N50 of 3801 for SAVAGE. For the 6-strain Poliovirus mixture we obtain similar results, yielding a major improvement of NGA50 values (1643 for SAVAGE compared to 7428 for Virus-VG) at the cost of a minor decrease in target haplotypes reconstructed (83.7% for SAVAGE compared to 80.7% for Virus-VG).

On both the ZIKV and Poliovirus data, we observe a slight increase in error rate after applying our method; however, Virus-VG leaves with an error rate of 0.115% (ZIKV) and 0.064% (Poliovirus), which is still extremely low. A thorough analysis turns up that this increase is due to errors in the input contigs that become more expressed only after having assembled the full-length haplotypes, so these errors are not primarily due to the method presented here. Moreover, the full-length contiguity of the haplotypes clearly offsets the minute shift in accuracy.

Finally, we analyze performance on a real benchmark, the labmix, and observe the same behaviour for Virus-VG: a significant improvement in NGA50 values (1450 for SAVAGE compared to 4642 for Virus-VG) but also an increase in error rate (0.066 for SAVAGE compared to 0.324 for Virus-VG). However, it is important to realize that the true sequences considered here may not fully represent the sample, because extremely high mutation rates allow the virus to mutate and recombine in vitro before sequencing.

### 3.4 Comparison with the state-of-the-art

Rows 3 and 4 in Tables 1a-1d display results for state-of-the-art methods PredictHaplo (Prabhakaran *et al*., 2014) and ShoRAH (Zagordi *et al*., 2011), run with default parameter settings. Both of these methods are reference-guided, hence cannot immediately be compared with Virus-VG, which operates entirely de novo. To simulate a de novo type scenario for these reference-guided approaches, we provided them with a bootstrap reference genome computed by running (Yang *et al*., 2012, VICUNA), a state-of-the-art tool for generating consensus virus genomes, on the input reads. We also ran (Ahn and Vikalo, 2018, aBayesQR) and (Chen *et al*., 2018, PEHaplo), but found them unsuitable for reconstructing full-length genomes at ultra deep coverage: they could not finish the job within 500 hours. Hence, we only present results for these methods on the HIV pol region data sets (both simulated and real) in Section 3.5.

We first evaluated both PredictHaplo and ShoRAH on our simulated data and, in all cases, we found our method to have (quite significant) advantages, in terms of accuracy, number of strains, and strain-specific genomes covered. As was already observed earlier Baaijens *et al*. (2017), reference-guided methods greatly depend on the quality of the reference genome provided and have to deal with biases towards the reference genome. This results in error rates which are 1.1–59 times higher than Virus-VG for PredictHaplo, and more than 12 times higher than Virus-VG for ShoRAH. At the same time, these methods miss a big fraction of the target haplotypes on all data sets except the labmix.

PredictHaplo and ShoRAH both had difficulty processing the Poliovirus data. A possible explanation is the high divergence between the virus strains and the reference genome used, leading to gaps in coverage when considering alignments to the reference genome, which tends to confuse reference-guided methods. In particular, two of the six strains have a big deletion (more than 1000bp) compared to both the reference genome and the other four strains; this may also explain the failure to run ShoRAH even using a bootstrap reference genome, as well as the extremely low target reconstructed for PredictHaplo. These results again highlight the advantage of a fully de novo approach compared to reference-guided methods.

### 3.5 Gene sequence reconstruction

Although our main goal is to reconstruct full length genomes, some studies also require sequence reconstruction from data corresponding to shorter regions of the genome. Therefore, we explored the ability of Virus-VG to reconstruct the HIV pol gene sequence, a 3kb region coding for DNA polymerase. This also allowed us to compare our assemblies against aBayesQR(Ahn and Vikalo, 2018) and PEHaplo (Chen *et al*., 2018), as these methods were unable to process full-length genomes at deep coverage.

Table 2 presents results for all methods on simulated data, the 7-strain HIV pol mixture, at sequencing depths of 500x, 1000x, and 5000x. This data set is very challenging, with pairwise divergence of only 1% and relative abundances ranging from 0.5% and 61.5%. Therefore, it is not surprising to see relatively low target reconstructed for all methods; the only method that is able to find the strain of 0.5% abundance is PEHaplo at 5000x coverage (see Supplementary Material). For all methods except ShoRAH, reconstructed target values improve upon increasing coverage. Virus-VG shows similar behaviour as on full-length genomes: it gives a major improvement in N50 and NGA50 values compared to SAVAGE, and while error rates are higher than for SAVAGE they remain much lower than for other methods.

**Table 2:** Assembly results for the simulated HIV pol region (∼3kb) at coverage 500x, 1000x and 5000x. ER = Error Rate, computed as the sum of the fraction of ‘N’s (ambiguous bases) and the mismatch- and indel rates. *If contigs are full-length, this number reflects the estimated number of strains in the quasispecies.

|  | # contigs* | target (%) | N50 | NGA50 | ER(%) |
| --- | --- | --- | --- | --- | --- |
| **500x** |  |  |  |  |  |
| SAVAGE | 7 | 51.0 | 2291 | 930 | 0.005 |
| Virus-VG | 4 | 52.7 | 2974 | 2330 | 0.057 |
| aBayesQR | 6 | 55.7 | 3069 | 3069 | 0.249 |
| PEHaplo | 64 | 55.7 | 3048 | 3062 | 0.459 |
| PredictHaplo | 1 | 14.3 | 3068 | - | 0.392 |
| ShoRAH | 27 | 61.0 | 2680 | 2680 | 1.322 |
| **1000x** |  |  |  |  |  |
| SAVAGE | 12 | 57.2 | 1416 | 986 | 0.012 |
| Virus-VG | 6 | 61.3 | 2977 | 2907 | 0.116 |
| aBayesQR | 6 | 62.8 | 3070 | 3070 | 0.258 |
| PEHaplo | 59 | 61.7 | 3045 | 3063 | 0.449 |
| PredictHaplo | 2 | 21.4 | 3070 | - | 0.514 |
| ShoRAH | 27 | 59.8 | 2680 | 2680 | 1.304 |
| **5000x** |  |  |  |  |  |
| SAVAGE | 17 | 66.4 | 1612 | 1596 | 0.005 |
| Virus-VG | 7 | 64.7 | 2853 | 2864 | 0.089 |
| aBayesQR | 7 | 61.4 | 3074 | 3074 | 0.283 |
| PEHaplo | 469 | 97.0 | 3045 | 3072 | 0.519 |
| PredictHaplo | 2 | 28.5 | 3070 | - | 0.587 |
| ShoRAH | 32 | 51.4 | 2766 | 2766 | 1.404 |

In addition, we selected reads from the pol region of the labmix (real data) and subsampled this selection to 100x and 1000x coverage, respectively. Again, Virus-VG achieves low error rates in combination with high N50/NGA50 values and improves as sequencing depth increases. To ensure robustness, simulations and subsampling were performed 10 times, methods were run on these 10 samples and results were averaged.

### 3.6 Haplotype abundance estimation

We also evaluated the accuracy of the abundance estimates obtained for each haplotype of the simulated data sets, since we know the exact true frequencies for each of the strains. The reconstructed sequences were aligned to the ground truth sequences and assigned to the closest matching strain. For each ground truth strain, we summed the abundance estimates of the sequences assigned to it, thus obtaining a total estimate for this strain. Then we compared this estimate to the true strain abundance and computed the absolute frequency estimation errors. In case of any missing strains, the true frequencies were normalized first, taking only the assembled sequences into account for a fair comparison. SAVAGE and PEHaplo assemblies are not evaluated as these methods do not provide abundance estimates.

Our method predicts highly accurate abundances for the reconstructed strains, with an average absolute estimation error of 0.1% on the HCV data, 0.3% on the ZIKV data, and 0.6% on the Poliovirus data. In comparison, PredictHaplo achieves an average absolute estimation error of 0.9% (HCV), 4.9% (ZIKV), and 10.6% (Polio), while ShoRAH is even further off with 8.5% (HCV) and 39% (ZIKV). On the HIV pol data, we observe higher estimation errors for all methods, with an absolute error of 2.6–5.8% for Virus-VG, 3.9–4.9% for aBayesQR, 8.2–22.1% for ShoRAH, and 6.4% for PredictHaplo (only evaluated at 5000x). Relative estimation errors show a similar pattern. A likely explanation for increased error rates on the HIV pol data is the complexity of the data set, with very low frequency strains in combination with a low total coverage (500x–5000x). A more detailed analysis can be found in the Supplementary Material.

Figure 2 shows the true haplotype frequencies versus the relative error^3^ per method. Results are clustered by true abundance into bins of size 0.05 and average errors are plotted for each bin. On the Poliovirus data there are no results for ShoRAH or PredictHaplo, because the first could not process this data set while the latter found less than a single strain. aBayesQR could only process the HIV pol data; on this data set, Virus-VG, PredictHaplo, and aBayesQR have a similar error pattern, with values well below the errors made by ShoRAH. On HCV and ZIKV data, however, we observe that Virus-VG outperforms the other methods in terms of frequency estimation, with estimates that are closest to the true values. An immediate interpretation of these findings is that accuracy in estimating abundance is inevitably linked with accuracy in haplotype reconstruction, which may explain our overall advantages.

**Figure 2:**
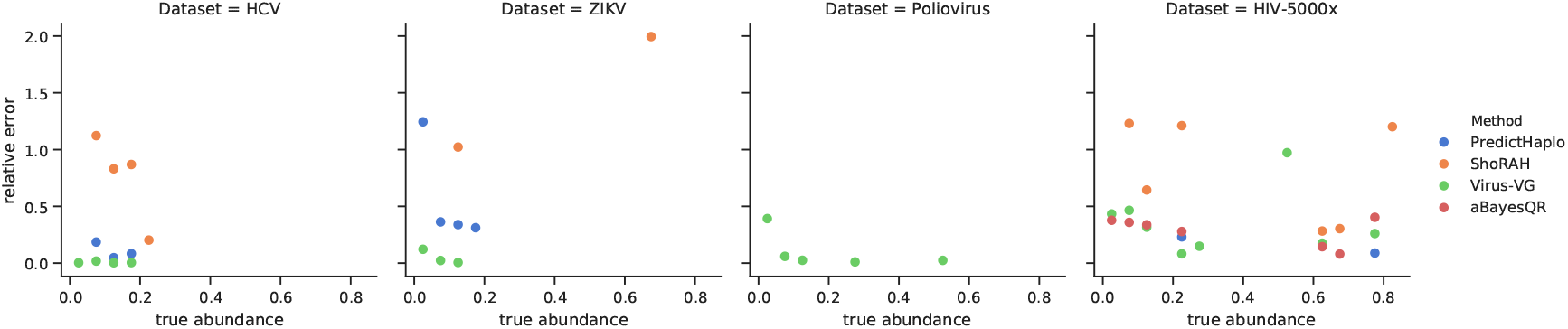
Relative errors for haplotype abundance estimation versus true strain frequencies. Results were evaluated per method, per data set, and binned by true frequency into bins of size 0.05. Plots show the average relative error per bin. True frequencies were normalized per assembly, taking only the assembled sequences into account for a fair comparison. Only assemblies containing at least 2 strains were evaluated. Plots for the simulated HIV data at 500x and 1000x are similar to HIV-5000x and presented in the Supplementary Material.

### 3.7 Runtime evaluation

By their worst-case runtime complexity, both candidate path generation and minimizing for selecting optimal sets of haplotypes reflect exponential procedures in Virus-VG. In practice, however, this is not an issue: on our benchmarks Virus-VG is 2.5–87 times faster than SAVAGE, which together form our de novo assembly pipeline. Combined, this pipeline takes 43–286 CPU hours on full-length data sets of 20,000x coverage. This is slower than PredictHaplo (2.0–7.4 CPU hours) but faster than ShoRAH (351–814 CPU hours), both of which are reference-guided. However, SAVAGE and Virus-VG can use multiple cores while PredictHaplo does not, leading to comparable wall clock times when multithreading. We present a more detailed runtime and memory analysis in the Supplementary Material.

### 3.8 Summary

We have benchmarked three de novo assembly tools (SAVAGE, Virus-VG, and PEHaplo) and three reference-guided methods (aBayesQR, PredictHaplo, and ShoRAH). The only methods that are stable with respect to all data sets considered, both full-length genomes and shorter regions, are SAVAGE and Virus-VG. Although SAVAGE achieves lowest error rates, Virus-VG is able to build full-length haplotypes with error rates slightly higher than SAVAGE, but still much lower than other methods. PredictHaplo performs well on the labmix at 20.000x, full-length genome and pol region, but it misses many haplotypes on all other data sets, with only 14–64% of the target genomes reconstructed. Virus-VG yields most accurate frequency estimates on full-length genomes, and performs similar to aBayesQR on the HIV pol region. In terms of CPU time, the combination of SAVAGE and Virus-VG is on a par with or faster than all other methods except PredictHaplo, which is consistently faster; in terms of wall clock time, however, both SAVAGE and Virus-VG achieve a major speedup by using multithreading, while PredictHaplo does not.

## 4 Discussion

We have presented an algorithm that turns viral strain-specific contigs, such as available from a de novo assembler like SAVAGE (Baaijens *et al*., 2017), into full-length, viral strain-specific haplotypes, without the use of a reference genome at any point. We first construct a contig variation graph, which arranges haplotype-specific contigs sampled from a viral quasispecies in a convenient and favorable manner. We then enumerate all maximal length paths through this graph that maximally concatenate the contig subpaths. Last, we solve a minimization problem that assigns abundance estimates to maximal length paths that are optimal in terms of being compatible with abundances computed for the nodes in the graph. We finally output the optimal such set of paths together with their abundances, by which *we have completed the de novo viral quasispecies assembly task*.

In benchmarking experiments, we have demonstrated that our method yields major improvements over the input contigs in terms of assembly length, while preserving high accuracy in terms of error rates. Compared to state-of-the-art viral quasispecies assemblers—all of which operate in a reference genome dependent manner—our method produces haplotype-resolved assemblies that are both more complete, in terms of haplotypes covered, and more accurate, in terms of error rates. We believe that (a) this reflects the strength of a fully *de novo* approach, because we avoid to deal with reference-induced biases. We also believe that (b) this is a result of directly integrating haplotype abundance estimation into reconstruction of haplotypes.

Still, improvements are possible. Our current optimization problem employs the absolute difference to determine the abundance estimation error. As future work, we consider the exploration of probabilistic error models, e.g., by modeling path abundance as being Poisson distributed (Medvedev *et al*., 2010) and calculating the likelihood of the observed node abundances.

Further, we had already alluded to that the number of candidate paths is exponential in the number of input contigs, which could theoretically be overwhelming when dealing with highly fragmented assembly output. Our runtime benchmarks show that this is not an issue with standard data sets. Nevertheless, we will consider more efficient alternative solutions in future work, based on a flow formulation of the problem that we recently found, yielding a yet to be implemented polynomial time algorithm.

## Funding

This work was supported by the Netherlands Organisation for Scientific Research (NWO) through Vidi grant 679.072.309, Veni grant 639.021.648 and Gravitation Programme Networks 024.002.003.

## Footnotes

1 https://github.com/jstjohn/SimSeq

2 https://github.com/cbg-ethz/5-virus-mix

3 | *x* − *x*\* | /(0.5(*x* + *x*\*))

