## Supplemental Material for "Full-length de novo viral quasispecies assembly through variation graph construction"

**6 Command lines** **21**

**7 Installation** **21**

[*]Centrum Wiskunde & Informatica, Amsterdam, Netherlands
[†]University Medical Center Utrecht, Utrecht, Netherlands
[‡]University of Duisberg-Essen, Essen, Germany
[§]Dana Farber Cancer Institute, Harvard Medical School, Boston, United States
[¶]Vrije Universiteit, Amsterdam, Netherlands
[‖]INRIA-Erable
[**]Utrecht University, Utrecht, Netherlands

### 1 Parameters

Our method requires to manually set three parameters, the minimal node abundance $\alpha$, the minimal haplotype abundance $\gamma$, and the maximal trim length $\tau$. The minimal node abundance $\alpha$ refers to removing mismatches when concatenating paths, see 'Correcting errors in paths $p \in P'$' in the main manuscript. As a general guideline, increasing $\alpha$ leads to increasing numbers of candidate paths, hence an increasing number of variables in the minimization problem. The greater the number of candidate paths, the greater the chance that the true haplotypes are present and while at the same time the greater the risk to pick up haplotype artifacts.

The minimal haplotype abundance $\gamma$ refers to selecting haplotypes after having solved the minimization problem in Section 2.2.2. Any haplotype $p \in P$ where $a(p) < \gamma$ will be discarded from the output. The reasoning for this threshold is that in general, de novo assemblers are unable to reconstruct contigs below a certain abundance threshold. Therefore, any haplotypes with an abundance below this threshold are likely the results of sequencing artifacts or assembly errors. Increasing $\gamma$ leads to a higher accuracy but also a loss of low abundance haplotypes; it is a common trade-off in quasispecies assembly. We recommend setting $\alpha$ and $\gamma$ to 0.5% and 1.0% of the total sequencing depth of the original data set, respectively. These default settings were chosen according to the quality of the input contigs (Baaijens *et al.*, 2017). Given that the full-length data sets considered here have a total sequencing depth of 20,000x, these experiments were run with $\alpha = 100$ and $\gamma = 200$. For data sets with lower coverage, $\alpha$ and $\gamma$ were set to 0.5% and 1.0% of the total sequencing depth sequencing depth, respectively.

The trim length $\tau$ refers to 'Trimming paths $p \in P'$' in Section 2.2.1. Due to issues in contig computation, uncorrected sequencing errors are often located on the extremities of the contig. Since contigs have large overlaps in general, we shorten the contig paths by removing their extremities for at most $\tau$ bases. Increasing $\tau$ leads to less concatenations of paths from $P'$, hence to less candidate paths in general, at the risk of not concatenating correctly joining contigs. Because of this risk, we recommend setting $\tau$ to small values only; its default value is 10bp and this value was used in all experiments.

### 2 Removing concatenations lacking physical evidence

We observed that during the de novo assembly process, coincidentally, situations like the one depicted in Figure 1 can occur. In this situation, the contig $q_2$ provides physical evidence for the concatenation of contigs $q_1 \rightarrow_c p_2$, while there is no such evidence for the concatenation $p_1 \rightarrow_c p_2$. At the same time, $p_1$ concatenates well with $q_4$, so we can safely remove $p_1 \rightarrow_c p2$ without turning $p_1$ into a dead end. This procedure reduces the number of spurious haplotypes, thus speeding up the candidate path generation and optimization steps.

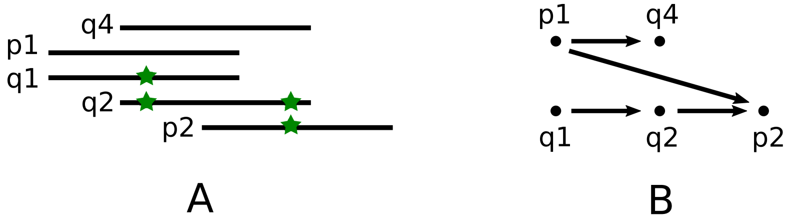

Figure 1: An illustration of concatenations lacking physical evidence. (A) Contigs $p_1, p_2, q_1, q_2, q_4$, with sequence variation indicated by a green star. (B) Possible contig concatenations are shown in the form of a graph: $p_1 \rightarrow_c q_4$, $q_1 \rightarrow_c p_2$, $q_1 \rightarrow_c q_2$, $q_2 \rightarrow_c p_2$. There is no contig $q_3$ in the assembly such that $p_1 \rightarrow_c q_3$ and $q_3 \rightarrow_c p_2$.

### 3 Data sets

#### 3.1 HCV and ZIKV

The HCV and ZIKV simulated data sets were presented by (Baaijens *et al.*, 2017) and are publicly available at `https://bitbucket.org/jbaaijens/savage-benchmarks`.

#### 3.2 Poliovirus

We extracted sequences for 6 closely related poliovirus strains from the NCBI nucleotide database, accession numbers MG212475.1, MG212489.1, MG212484.1, MG21469.1, MG212490.1, and MG212491.1. Two of these sequences (MG212476.1 and MG21484.1) show a big deletion (larger than 1000bp) compared to the Sabin2 reference strain. We used SimSeq[1] to simulate 2x250bp Illumina Miseq reads (using the MiSeq error profiles) – see Section 6. Strain frequencies increase exponentially (1.6%, 3.2%, 6.3%, 12.7%, 25.4%, 50.8%) with a total coverage of 20.000x.

#### 3.3 HIV pol region

We downloaded a sequence for the HIV-1 pol region from the NCBI nucleotide database (accession number D86068.1) and generated 7 mutated strains by introducing random mutations (mostly substitutions, some small indels) at a rate of 0.5%. Thus, each pair of strains has a pairwise divergence of approximately 1%. For these 7 strains we simulated 2x250bp Illumina Miseq reads using SimSeq, with relative strain frequencies of 0.5%, 1%, 2%, 5%, 10%, 20%, 61.5%. We built 3 such data sets with varying total coverage: 500x, 1000x, and 5000x, respectively. For each of these data sets, simulations were performed 10 times and assembly tools were run on all 10 samples to ensure robustness; average results are presented for each data set.

#### 3.4 Labmix

The labmix data set consists of 5 well-known HIV-1 strains that were mixed in a lab and then sequenced using Illumina MiSeq technology (Di Giallonardo *et al.*, 2014). This is real sequencing data, so before further processing we the raw reads were trimmed using CutAdapt (Martin, 2011) (removing adapter sequences) and we checked for overlaps between forward and reverse reads using PEAR (Zhang *et al.*, 2014) – see Section 6 for the corresponding shell commands. The overall sequencing depth of this data set is approximately 20.000x.

###### Long terminal repeats

The HIV-1 sequence has repeats on the suffix and prefix of its genome, the so-called Long Terminal Repeats (LTR). When constructing a "consensus reference" from the labmix data set to use for reference-guided assembly tools, VICUNA (Yang *et al.*, 2012) gets confused by the repeats (LTR) and produces a reference genome that is much longer than the HIV genome. This, of course, has a great impact on the assemblies obtained with reference-guided methods, as these depend highly on the quality of the given reference genome. Moreover, many methods assume genomes to be repeat-free, or to have only small repeats (shorter than the read length); see e.g. Malhotra *et al.* (2016), Yang *et al.* (2012), Di Giallonardo *et al.* (2014), Astrovskaya *et al.* (2011). Therefore, we decided to benchmark on the labmix after excluding the terminal repeats, similar to Di Giallonardo *et al.* (2014).

In order to exclude the terminal repeats, we aligned the data to the HIV-1 reference sequence and removed all reads mapping to the long terminal repeats.

[1] `https://github.com/jstjohn/SimSeq`

##### Pol region

For studying assembly quality on shorter regions, we extracted the HIV pol region from the labmix data. We aligned all reads to the ground truth haplotypes, and selected reads belonging to the pol gene for each haplotype using samtools. In addition, to evaluate assembly quality on lower coverage data sets, we applied subsampling using the samtools -s option. We created data sets of 100x and 1000x coverage, 10 samples each to ensure robustness. Methods were run on all 10 samples and average results are presented for each data set.

### 4 Runtime and memory usage

#### 4.1 Detailed Virus-VG runtimes

The Virus-VG pipeline consists of two major steps, each of which can be divided into multiple smaller steps. First, we have graph construction, which requires contig alignment (MSA), graph indexing, and read mapping. Then, we have haplotype reconstruction, which consists of candidate path generation and path selection through optimization. We analyze Virus-VG runtimes for each of these steps to achieve a better understanding of the algorithmic bottlenecks. We also report the sizes (number of nodes and edges) of the corresponding variation graphs, as well as the number of candidate paths. All numbers are presented in Tables 1 and 2.

We observe that in general read mapping is the most expensive step in our workflow. However, as the assemblies become more complex, the number of candidate paths increases and hence candidate path generation becomes quite expensive. In practice, the complete Virus-VG workflow is still much faster than the de novo assembly step with SAVAGE (see section 4.2).

#### 4.2 Comparison to the state-of-the-art

We present runtime results (CPU hours and wall clock time) and peak memory usage for all methods on all data sets in Tables 3–5. Some methods (SAVAGE, Virus-VG, PEHaplo, and ShoRAH) can use multithreading; these methods were allowed to use 8 threads. All experiments were performed on a 24-core (Intel-Xeon 2.0GHz) Linux machine.

| | HCV | ZIKV | Poliovirus |
|---|---|---|---|
| graph construction | 21182 | 44894 | 12569 |
| *contig alignment* | 112 | 577 | 133 |
| *graph indexing* | 1.0 | 2.2 | 0.9 |
| *read mapping* | 19222 | 41930 | 11480 |
| haplotype reconstruction | 42 | 133 | 224 |
| *candidate path generation* | 14 | 60 | 200 |
| *path selection* | 9.5 | 26 | 13 |
| # nodes | 4547 | 6667 | 2065 |
| # edges | 6192 | 9179 | 2785 |
| # candidate paths | 17 | 93 | 889 |

Table 1: Detailed runtimes (CPU seconds) and graph statistics for Virus-VG on simulated full-length viral quasispecies data.

| | simulated HIV pol | | | real HIV pol (labmix) | | |
|---|---|---|---|---|---|---|
| | 500x | 1000x | 5000x | 100x | 1000x | 20.000x |
| graph construction | 52 | 113 | 576 | 170 | 517 | 19212 |
| *contig alignment* | 4.5 | 7.0 | 14 | 38 | 89 | 820 |
| *graph indexing* | 0.6 | 0.6 | 0.7 | 1.3 | 2.0 | 6.3 |
| *read mapping* | 36 | 87 | 507 | 118 | 383 | 17942 |
| haplotype reconstruction | 4.1 | 3.1 | 3.3 | 6.5 | 14 | 33843 |
| *candidate path generation* | 0.2 | 0.3 | 0.6 | 0.4 | 2.3 | 33216 |
| *path selection* | 0.1 | 0.1 | 0.1 | 0.4 | 0.7 | 329 |
| # nodes | 139 | 166 | 181 | 374 | 677 | 731 |
| # edges | 187 | 223 | 244 | 494 | 911 | 985 |
| # candidate paths | 5 | 8 | 12 | 23 | 51 | 65558 |

Table 2: Detailed runtimes (CPU seconds) and graph statistics for Virus-VG on simulated and real data for the HIV pol region.

| | CPU hours | Wall time | Peak memory usage (GB) |
|---|---|---|---|
| **HCV** | | | |
| SAVAGE | 38.6 | 6h19 | 26 |
| Virus-VG | 5.9 | 1h00 | 0.9 |
| aBayesQR | - | > 500h | - |
| PEHaplo | - | - | - |
| PredictHaplo | 2.7 | 2h43 | 1.1 |
| ShoRAH | 509 | 56h48 | 8.9 |
| **ZIKV** | | | |
| SAVAGE | 30.6 | 5h13 | 13 |
| Virus-VG | 12.5 | 1h17 | 0.8 |
| aBayesQR | - | > 500h | - |
| PEHaplo | - | - | - |
| PredictHaplo | 7.4 | 7h40 | 1.1 |
| ShoRAH | 814 | 104h35 | 10 |
| **Polio** | | | |
| SAVAGE | 60.2 | 10h04 | 3.3 |
| Virus-VG | 3.6 | 40m | 0.6 |
| aBayesQR | - | > 500h | - |
| PEHaplo | - | > 500h | - |
| PredictHaplo | 2.0 | 2h00 | 0.8 |
| ShoRAH | - | - | - |

Table 3: Runtime and -space comparison on simulated data sets for full length viral genomes (HCV, ZIKV, Polio) at 20.000x coverage. PEHaplo crashed on HCV and ZIKV, ShoRAH crashed on Polio and aBayesQR could not finish within 500h on any of these data sets.

| | CPU minutes | Wall time | Peak memory usage (GB) |
|---|---|---|---|
| **HIV pol 500x** | | | |
| SAVAGE | 15 | 2m52 | 1.1 |
| Virus-VG | 0.9 | 35s | 0.1 |
| aBayesQR | 1.5 | 2m41 | 0.1 |
| PEHaplo | 1.8 | 1m18 | 0.5 |
| PredictHaplo | 0.5 | 27s | 0.01 |
| ShoRAH | 26 | 4m42 | 0.07 |
| **HIV pol 1000x** | | | |
| SAVAGE | 144 | 18m00 | 2.3 |
| Virus-VG | 1.9 | 1m03 | 0.1 |
| aBayesQR | 0.9 | 1m16 | 0.2 |
| PEHaplo | 38 | 7m54 | 0.9 |
| PredictHaplo | 2.8 | 2m50 | 0.03 |
| ShoRAH | 95 | 21m55 | 0.1 |
| **HIV pol 5000x** | | | |
| SAVAGE | 849 | 2h42 | 2.6 |
| Virus-VG | 9.7 | 2m34 | 0.2 |
| aBayesQR | 27 | 43m47 | 1.0 |
| PEHaplo | 11869 | 27h37 | 12.4 |
| PredictHaplo | 3.5 | 3m31 | 0.1 |
| ShoRAH | 1749 | 3h44 | 0.5 |

Table 4: Runtime and -space comparison on simulated data for the HIV pol region ($\sim$3kb) at a coverage of 500x, 1000x, and 5000x, respectively.

| | CPU hours | Wall time | Peak memory usage (GB) |
|---|---|---|---|
| **Labmix pol 100x** | | | |
| SAVAGE | 0.01 | 15s | 0.4 |
| Virus-VG | 0.05 | 1m47 | 0.1 |
| aBayesQR | 1.94 | 1h58 | 0.2 |
| PEHaplo | 0.02 | 45s | 0.08 |
| PredictHaplo | 0.02 | 1m24 | 0.02 |
| ShoRAH | 0.10 | 37s | 0.04 |
| **Labmix pol 1000x** | | | |
| SAVAGE | 2.3 | 16m49 | 2.4 |
| Virus-VG | 0.1 | 3m27 | 0.2 |
| aBayesQR | 1.7 | 1h45 | 2.1 |
| PEHaplo | 50 | 7h59 | 10.8 |
| PredictHaplo | 0.05 | 3m06 | 0.03 |
| ShoRAH | 5.3 | 33m02 | 0.3 |
| **Labmix pol 20.000x** | | | |
| SAVAGE | 97 | 10h56 | 3.9 |
| Virus-VG | 15 | 12h27 | 6.3 |
| aBayesQR | 411 | 488h06 | 10.3 |
| PEHaplo | 6.6 | 2h58 | 8.2 |
| PredictHaplo | 0.9 | 52m34 | 0.5 |
| ShoRAH | 192 | 22h41 | 8.5 |
| **Labmix* 20.000x** | | | |
| SAVAGE | 276 | 31h34 | 7.2 |
| Virus-VG | 10 | 2h19 | 0.6 |
| aBayesQR | - | $> 500$h | - |
| PEHaplo | - | $> 500$h | - |
| PredictHaplo | 4.9 | 5h00 | 1.0 |
| ShoRAH | 351 | 40h36 | 10.7 |

Table 5: Runtime and -space comparison on real data (labmix) at various sequencing depths (100x, 1000x, 20.000x). The HIV pol-region constitutes approximately 3kb. *Full genome excluding long terminal repeats (LTR), constitutes approximately 9kb.

#### 5 Detailed assembly statistics

All assembly statistics per data set for each of the methods (SAVAGE, Virus-VG, aBayesQR, PEHaplo, PredictHaplo, and ShoRAH) can be found in Tables 6–9. Assembly statistics were computed using QUAST (Gurevich *et al.*, 2013). To cite the QUAST manual (`http://quast.bioinf.spbau.ru/manual.html`), these measures are computed as follows:

> **Genome fraction (%)** is the percentage of aligned bases in the reference genome. A base in the reference genome is aligned if there is at least one contig with at least one alignment to this base. Contigs from repetitive regions may map to multiple places, and thus may be counted multiple times.
> **N50** is the length for which the collection of all contigs of that length or longer covers at least half an assembly.
> **NG50** is the length for which the collection of all contigs of that length or longer covers at least half the reference genome.
> **NA50, NGA50** ("A" stands for "aligned") are similar to the corresponding metrics without "A", but in this case aligned blocks instead of contigs are considered. Aligned blocks are obtained by breaking contigs at misassembly events and removing all unaligned bases.
> **# N's per 100 kbp** is the average number of uncalled bases (N's) per 100000 assembly bases.
> **# mismatches per 100 kbp** is the average number of mismatches per 100000 aligned bases.
> **# indels per 100 kbp** is the average number of indels per 100000 aligned bases. Several consecutive single nucleotide indels are counted as one indel.

We refer to "Genome fraction (%)" as "target (%)" and by "Error Rate" we refer to the overall error rate, which equals the sum of N-rate, mismatch rate, and indel rate. This measure reflects how much one can trust a given contig. False discovery affects error rate and target (%) if contigs can be aligned, which is the case here. In other words, incorrect contigs do lead to a higher target (%), but also to an increased error rate. Together this gives an impression of assembly accuracy.

Tables 9 and 8 show results on simulated and real data, respectively, for the HIV pol region ($\sim$ 3kb). Note that the primary intention of Virus-VG is to reconstruct full-length genomes and the method, by its nature, profits from full-length genomes. Reconstructing isolated regions is not necessarily the point for a de novo approach; if isolated well-known regions are addressed, reference-based methods might be a good option. However, it is interesting here because it allows us to compare also to aBayesQR and PEHaplo.

For the labmix pol data at 100x coverage, we observe that SAVAGE can only reconstruct 9.8% of the target genomes (Table 8. Since these contigs are used as input for Virus-VG, our method cannot be expected to yield a complete assembly, but target coverage does increase to 21.9% after applying Virus-VG.

We observe that PredictHaplo performs well on the labmix at 20.000x, full-length genome and pol region, but it misses many haplotypes on all other data sets, with only 14–64% of the target genomes reconstructed. It seems that PredictHaplo was overfitted towards the 20.000x labmix data; in fact, this data set comes together with the PredictHaplo software. Even on the labmix data at lower coverage of 100x and 1000x, PredictHaplo reconstructs only 1 and 3 strains, respectively.

| | # contigs* | target (%) | N50 | NGA50 | ER(%) | recall | precision |
|---|---|---|---|---|---|---|---|
| SAVAGE | 26 | 99.4 | 8964 | 8964 | 0.001 | 1.0 | 0.92 |
| Virus-VG | 10 | 99.3 | 9281 | 9203 | 0.001 | 0.90 | 0.90 |
| PredictHaplo | 9 | 73.8 | 7636 | 7608 | 0.059 | 0.10 | 0.11 |
| ShoRAH | 639 | 56.9 | 7570 | 7570 | 4.294 | 0 | 0 |

(a) 10-strain HCV mixture (simulated Illumina MiSeq)

| | # contigs* | target (%) | N50 | NGA50 | ER(%) | recall | precision |
|---|---|---|---|---|---|---|---|
| SAVAGE | 100 | 98.8 | 2954 | 3801 | 0.023 | 1.0 | 0.77 |
| Virus-VG | 20 | 92.8 | 10202 | 10210 | 0.115 | 0.53 | 0.40 |
| PredictHaplo | 8 | 53.3 | 10270 | 10267 | 0.126 | 0.20 | 0.38 |
| ShoRAH | 493 | 26.3 | 10117 | 10117 | 4.392 | 0 | 0 |

(b) 15-strain ZIKV mixture (simulated Illumina MiSeq)

| | # contigs* | target (%) | N50 | NGA50 | ER(%) | recall | precision |
|---|---|---|---|---|---|---|---|
| SAVAGE | 59 | 83.7 | 1089 | 1643 | 0.019 | 1.0 | 0.88 |
| Virus-VG | 14 | 80.7 | 7316 | 7428 | 0.064 | 0.17 | 0.10 |
| PredictHaplo | 3 | 16.6 | 7461 | - | 1.825 | 0 | 0 |

(c) 6-strain Poliovirus mixture (simulated Illumina MiSeq)

Table 6: Assembly results for simulated deep sequencing data. ER = Error Rate, computed as the sum of the fraction of 'N's (ambiguous bases) and the mismatch- and indel rates. ShoRAH could not process the Poliovirus data. PEHaplo and aBayesQR could not process any of these data sets. *If contigs are full-length, this number reflects the estimated number of strains in the quasispecies.

| | # contigs* | target (%) | N50 | NGA50 | ER(%) | recall | precision |
|---|---|---|---|---|---|---|---|
| SAVAGE | 68 | 97.9 | 1026 | 1450 | 0.066 | 1.0 | 0.25 |
| Virus-VG | 23 | 90.6 | 2130 | 4642 | 0.324 | 0.80 | 0.22 |
| PredictHaplo | 6 | 100.0 | 8825 | 8825 | 1.066 | 0 | 0 |
| ShoRAH | 250 | 100.0 | 8775 | 8775 | 3.910 | 0 | 0 |

Table 7: Labmix whole genome (excluding LTR) at full coverage (20.000x). ER = Error Rate, computed as the sum of the fraction of 'N's (ambiguous bases) and the mismatch- and indel rates. Note that PEHaplo and aBayesQR could not process this data set. *If contigs are full-length, this number reflects the estimated number of strains in the quasispecies.

| | # contigs* | target (%) | N50 | NGA50 | ER(%) | recall | precision |
|---|---|---|---|---|---|---|---|
| **100x coverage** | | | | | | | |
| SAVAGE | 2 | 9.8 | 667 | - | 0.039 | 1.000 | 0.848 |
| Virus-VG | 5 | 21.9 | 924 | - | 0.478 | 0.960 | 0.586 |
| aBayesQR | 8 | 95.4 | 3454 | 3000 | 1.220 | 0.000 | 0.000 |
| PEHaplo | 6 | 46.3 | 2869 | - | 1.180 | 0.000 | 0.000 |
| PredictHaplo | 1 | 19.8 | 3450 | - | 1.490 | 0.000 | 0.000 |
| ShoRAH | 52 | 99.2 | 3406 | 3000 | 1.533 | 0.000 | 0.000 |
| **1000x coverage** | | | | | | | |
| SAVAGE | 15 | 91.2 | 1378 | 1420 | 0.063 | 1.000 | 0.243 |
| Virus-VG | 9 | 93.0 | 3071 | 2888 | 0.297 | 0.400 | 0.151 |
| aBayesQR | 7 | 84.0 | 3520 | 3023 | 1.332 | 0.000 | 0.000 |
| PEHaplo | 358 | 99.8 | 3488 | 3023 | 1.768 | 0.060 | 0.001 |
| PredictHaplo | 3 | 64.0 | 3517 | 3023 | 0.611 | 0.000 | 0.000 |
| ShoRAH | 122 | 100.0 | 3474 | 3023 | 1.825 | 0.000 | 0.000 |
| **20.000x coverage** | | | | | | | |
| SAVAGE | 45 | 95.5 | 704 | 897 | 0.014 | 1.000 | 0.270 |
| Virus-VG | 32 | 95.4 | 1674 | 3011 | 0.770 | 0.200 | 0.030 |
| aBayesQR | 7 | 40.0 | 3522 | 3023 | 1.760 | 0.000 | 0.000 |
| PEHaplo | 1667 | 100.0 | 1620 | 3023 | 1.979 | 0.000 | 0.000 |
| PredictHaplo | 5 | 100.0 | 3523 | 3023 | 0.276 | 0.000 | 0.000 |
| ShoRAH | 200 | 100.0 | 3484 | 3023 | 1.661 | 0.000 | 0.000 |

Table 8: Labmix pol region at coverage 100x, 1000x and 20.000x. ER = Error Rate, computed as the sum of the fraction of 'N's (ambiguous bases) and the mismatch- and indel rates. Note that reconstructing isolated regions is not the primary intention of Virus-VG, as it is a de novo approach. *If contigs are full-length, this number reflects the estimated number of strains in the quasispecies.

| | # contigs* | target (%) | N50 | NGA50 | ER(%) | recall | precision |
|---|---|---|---|---|---|---|---|
| **500x** | | | | | | | |
| SAVAGE | 7 | 51.0 | 2291 | 930 | 0.005 | 0.584 | 0.941 |
| Virus-VG | 4 | 52.7 | 2974 | 2330 | 0.057 | 0.416 | 0.660 |
| aBayesQR | 6 | 55.7 | 3069 | 3069 | 0.249 | 0.000 | 0.000 |
| PEHaplo | 64 | 55.7 | 3048 | 3062 | 0.459 | 0.000 | 0.000 |
| PredictHaplo | 1 | 14.3 | 3068 | - | 0.392 | 0.000 | 0.000 |
| ShoRAH | 27 | 61.0 | 2680 | 2680 | 1.322 | 0.000 | 0.000 |
| **1000x** | | | | | | | |
| SAVAGE | 12 | 57.2 | 1416 | 986 | 0.012 | 0.710 | 0.966 |
| Virus-VG | 6 | 61.3 | 2977 | 2907 | 0.116 | 0.400 | 0.443 |
| aBayesQR | 6 | 62.8 | 3070 | 3070 | 0.258 | 0.000 | 0.000 |
| PEHaplo | 59 | 61.7 | 3045 | 3063 | 0.449 | 0.014 | 0.004 |
| PredictHaplo | 2 | 21.4 | 3070 | - | 0.514 | 0.000 | 0.000 |
| ShoRAH | 27 | 59.8 | 2680 | 2680 | 1.304 | 0.000 | 0.000 |
| **5000x** | | | | | | | |
| SAVAGE | 17 | 66.4 | 1612 | 1596 | 0.005 | 0.755 | 0.937 |
| Virus-VG | 7 | 64.7 | 2853 | 2864 | 0.089 | 0.528 | 0.579 |
| aBayesQR | 7 | 61.4 | 3074 | 3074 | 0.283 | 0.000 | 0.000 |
| PEHaplo | 469 | 97.0 | 3045 | 3072 | 0.519 | 0.000 | 0.000 |
| PredictHaplo | 2 | 28.5 | 3070 | - | 0.587 | 0.000 | 0.000 |
| ShoRAH | 32 | 51.4 | 2766 | 2766 | 1.404 | 0.000 | 0.000 |

Table 9: Simulated HIV pol region at coverage 500x, 1000x and 5000x. ER = Error Rate, computed as the sum of the fraction of 'N's (ambiguous bases) and the mismatch- and indel rates. Note that reconstructing isolated regions is not the primary intention of Virus-VG, as it is a de novo approach. *If contigs are full-length, this number reflects the estimated number of strains in the quasispecies.

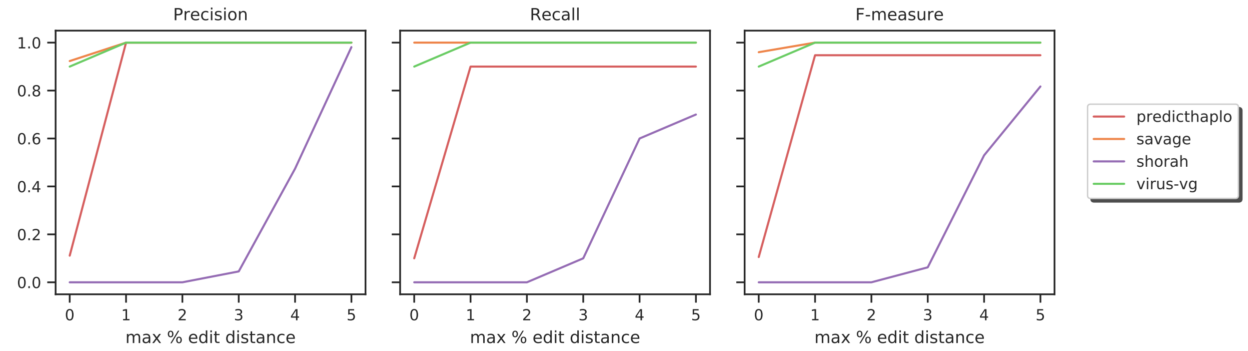

Figure 2: 10-strain HCV mixture (simulated Illumina MiSeq).

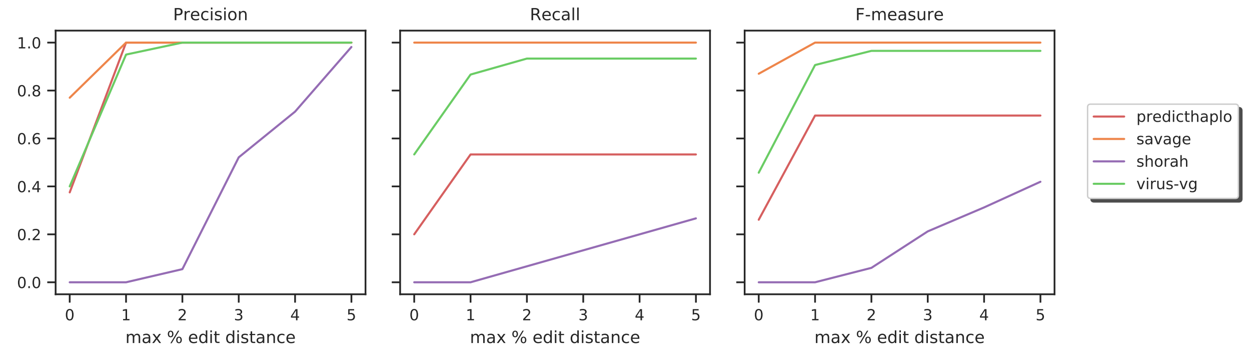

Figure 3: 15-strain ZIKV mixture (simulated Illumina MiSeq).

##### 5.1 Recall and precision

In addition to the QUAST assembly statistics, Tables 6–9 also present precision and recall (a.k.a. sensitivity and positive predictive value). We define recall as the number of true haplotypes having a contig that aligns with edit distance 0, divided by the total number of true haplotypes. We define precision as the number of contigs that align to a true haplotype with edit distance 0, divided by the total number of contigs.

These measures give SAVAGE a big advantage, because with short contigs it achieves much more exactly matching haplotypes (true positives). Virus-VG also achieves non-zero recall and precision on all data sets, but the other methods produce hardly any true positives: only PredictHaplo (HCV, ZIKV) and PEHaplo (simulated HIV pol 1000x, labmix pol 1000x) are able to achieve some true positives. We provide more detailed results on precision and recall in Figures 2–8. Here, we consider various thresholds for the relative edit distance (i.e. edit distance divided by alignment length) for contigs to be considered true positives. In addition, we also plot the F-measure (2*precision*recall/(precision+recall)). We observe that in general, SAVAGE achieves best results on these three measures, with high values for recall and precision already at low relative edit distance. However, SAVAGE only assembles short contigs. Virus-VG outperforms all other methods on all data sets, except for PEHaplo on the simulated HIV pol mixture at 5000x coverage. Note that in this case, also SAVAGE is outperformed by PEHaplo. On the labmix pol region at full coverage (20.000x), PredictHaplo achieves higher precision and F-measure when an edit distance of at least 1% is allowed for contigs to be considered true positives.

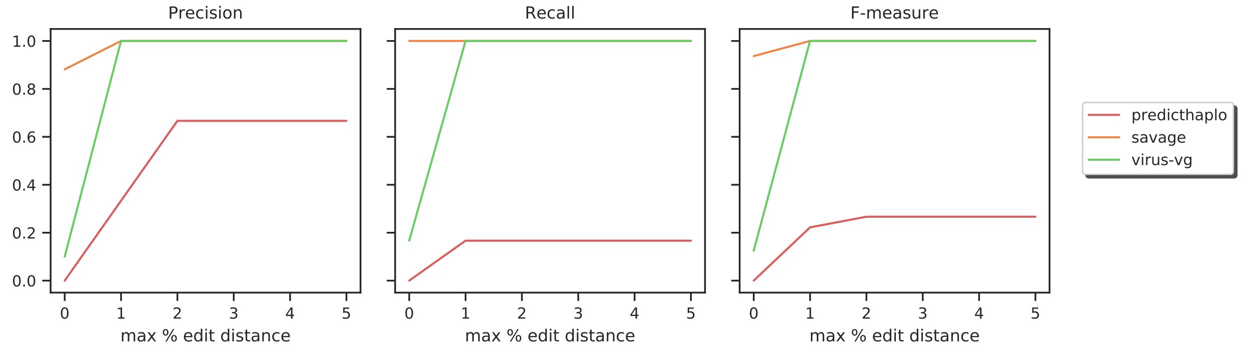

Figure 4: 6-strain Poliovirus mixture (simulated Illumina MiSeq).

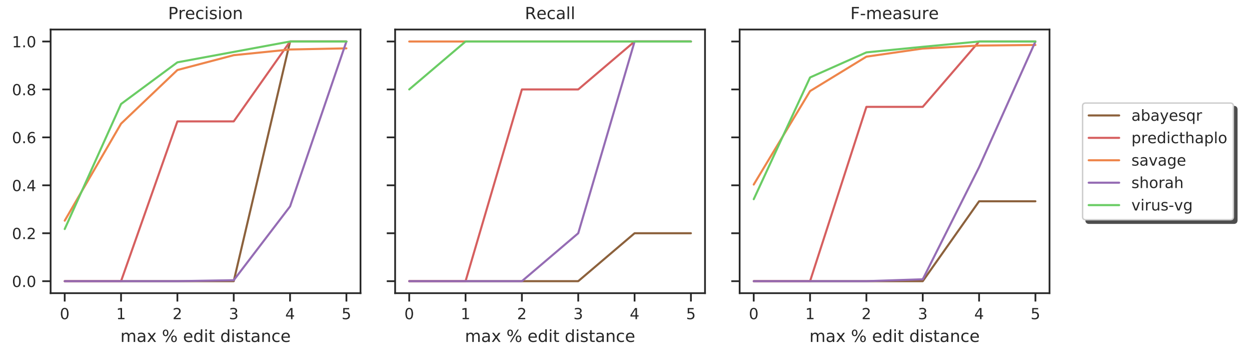

Figure 5: Labmix whole genome (excluding LTR) at full coverage (20.000x).

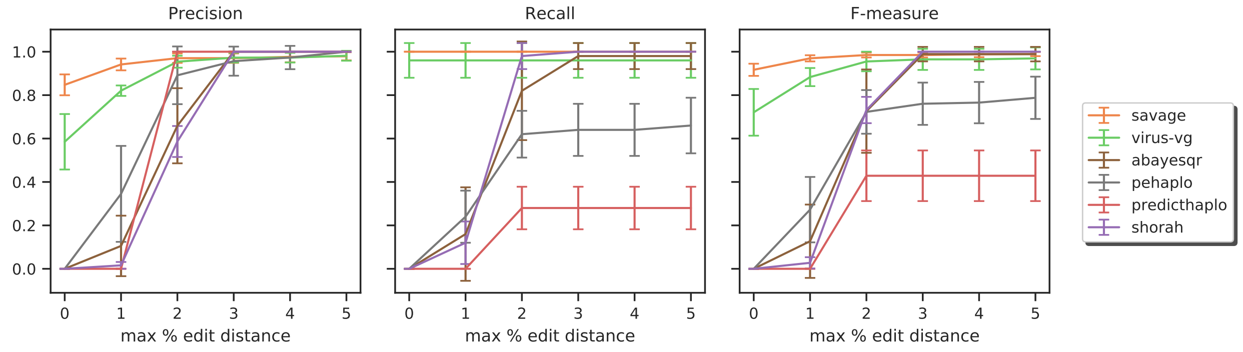

Figure 6: Labmix pol region subsampled at 100x coverage (average results over 10 subsamples, error bars indicate standard deviation).

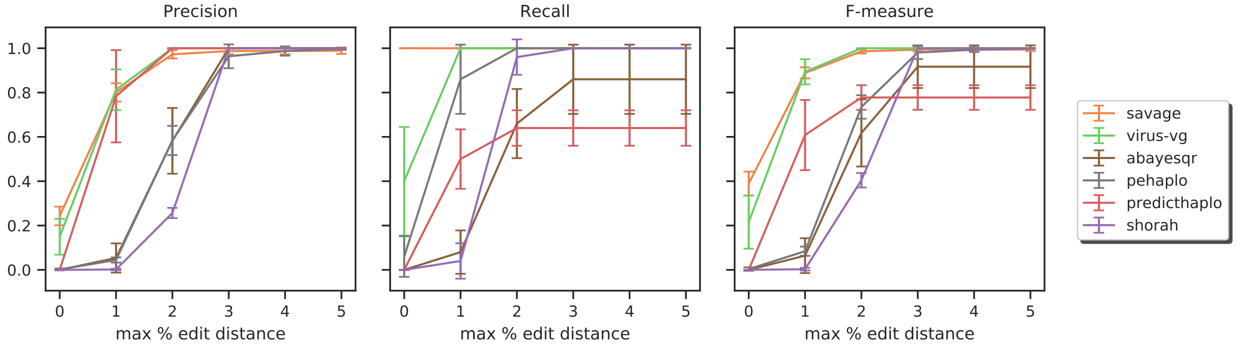

Figure 7: Labmix pol region subsampled at 1000x coverage (average results over 10 subsamples, error bars indicate standard deviation).

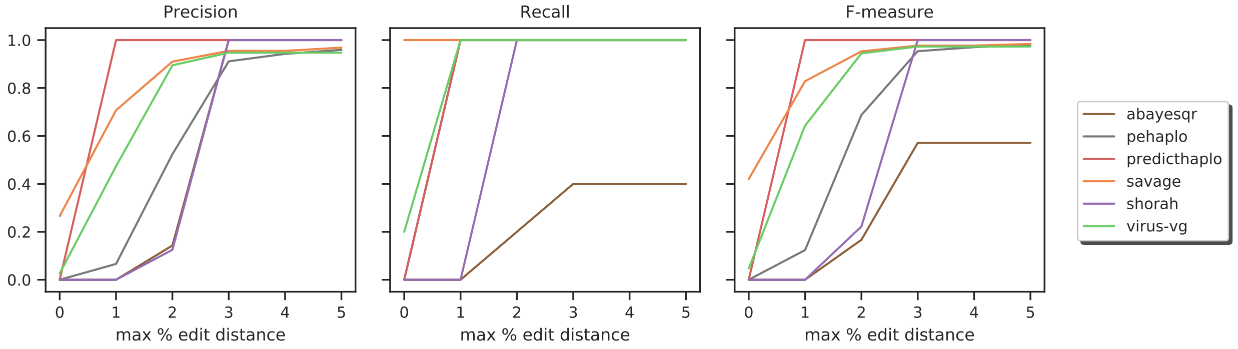

Figure 8: Labmix pol region at full coverage (20.000x).

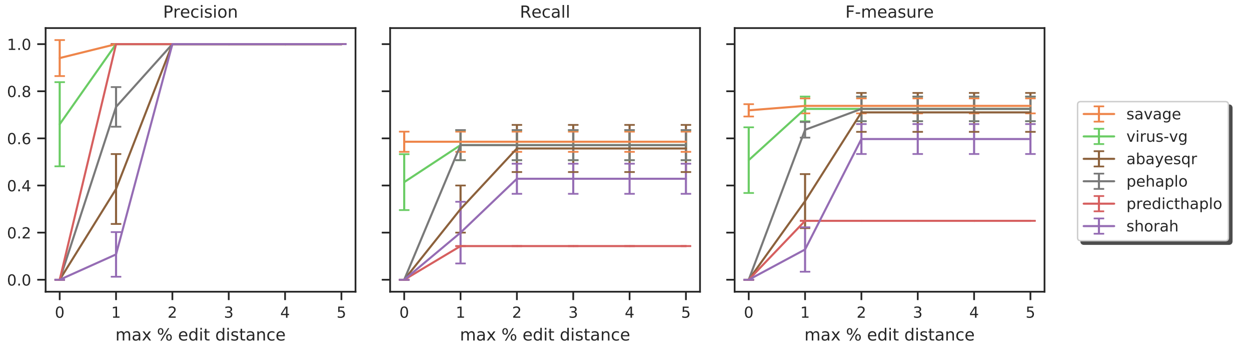

Figure 9: Simulated 7-strain HIV pol mixture at 500x coverage (average results over 10 simulations, error bars indicate standard deviation)

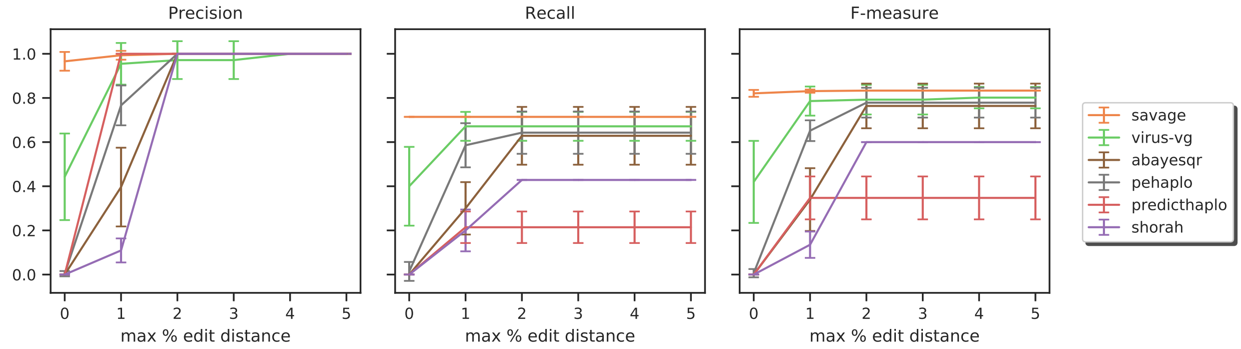

Figure 10: Simulated 7-strain HIV pol mixture at 1000x coverage (average results over 10 simulations, error bars indicate standard deviation)

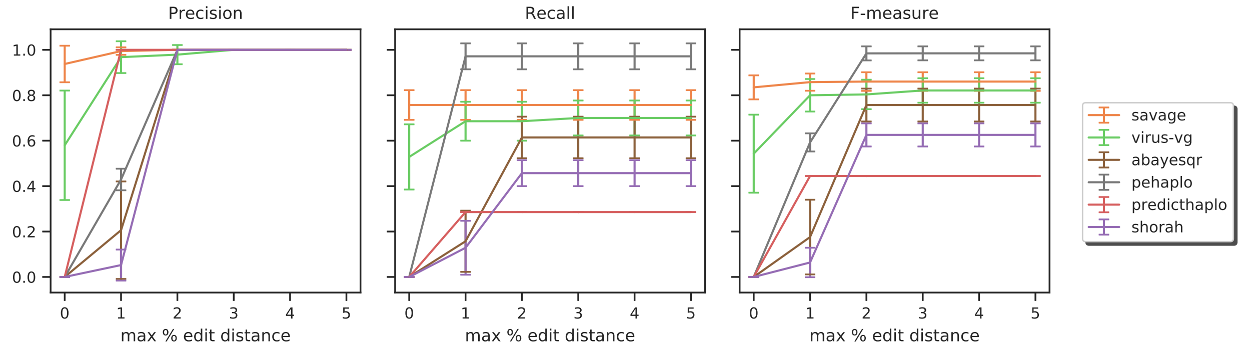

Figure 11: Simulated 7-strain HIV pol mixture at 5000x coverage (average results over 10 simulations, error bars indicate standard deviation)

#### 5.2 Frequency estimation

Abundance estimation errors per method are presented in Tables 10 and 11. Note that PEHaplo did not provide abundance estimates. Figure 12 shows scatterplots for all simulated data sets, evaluating relative abundance estimation errors versus relative strain abundance. Results are binned by strain abundance into bins of size 0.05 and average errors per bin are shown.

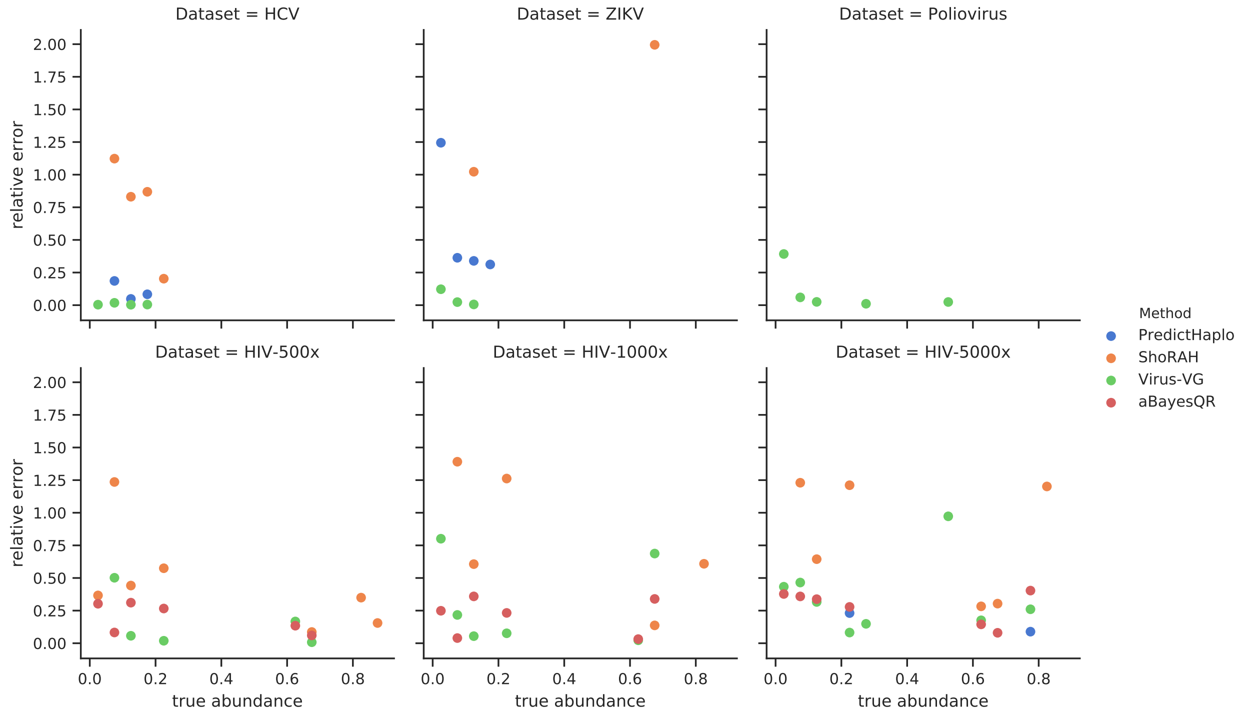

Figure 12: Relative errors for haplotype abundance estimation versus true strain frequencies. Results were evaluated per method, per data set, and binned by true frequency into bins of size 0.05. Plots show the average relative error per bin. True frequencies were normalized per assembly, taking only the assembled sequences into account for a fair comparison. Only assemblies containing at least 2 strains were evaluated.

#### 5.3 Strain level results

For each data set, for each assembly, we calculated the relative edit distance (edit distance / contig length) for each true haplotype to the closest reconstructed haplotype; results are presented in Tables 12–19.

| | HCV | | ZIKV | | Poliovirus | |
|---|---|---|---|---|---|---|
| | absolute error (%) | relative error (%) | absolute error (%) | relative error (%) | absolute error (%) | relative error (%) |
| Virus-VG | 0.1 | 0.9 | 0.3 | 6.0 | 0.6 | 12.8 |
| PredictHaplo | 0.9 | 11.3 | 4.9 | 69 | - | - |
| ShoRAH | 8.5 | 64 | 39 | 229 | - | - |

Table 10: Absolute and relative abundance estimation errors per method. For each data set, we present the average error over all assembled strains. Note that ShoRAH was unable to process the Poliovirus data, aBayesQR could not process any of these data sets, and PredictHaplo only found one of the six virus strains in the Poliovirus data set.

| | 500x | | 1000x | | 5000x | |
|---|---|---|---|---|---|---|
| | absolute error (%) | relative error (%) | absolute error (%) | relative error (%) | absolute error (%) | relative error (%) |
| Virus-VG | 3.2 | 32.2 | 2.6 | 43.6 | 5.8 | 76.0 |
| aBayesQR | 3.9 | 24.8 | 4.0 | 26.9 | 4.9 | 37.3 |
| PredictHaplo | - | - | - | - | 6.4 | 17.3 |
| ShoRAH | 8.2 | 48.2 | 13.2 | 75.8 | 22.1 | 139 |

Table 11: Absolute and relative abundance estimation errors per method on the simulated data for the HIV pol region at various sequencing depths. For each data set, we present the average error over all assembled strains.

| | s1 | s2 | s3 | s4 | s5 | s6 | s7 | s8 | s9 | s10 |
|---|---|---|---|---|---|---|---|---|---|---|
| *abundance* | 12% | 5% | 8% | 12% | 10% | 5% | 13% | 10% | 6% | 19% |
| SAVAGE | 0.0 | 0.0 | 0.0 | 0.0 | 0.0 | 0.0 | 0.0 | 0.0 | 0.0 | 0.0 |
| Virus-VG | 0.0 | 0.0 | 0.0 | 0.0 | 0.0 | 0.0 | 0.0 | 0.0 | 0.0 | 0.0 |
| PredictHaplo | 0.0 | 0.0 | 0.0 | 0.0 | 0.0 | - | 0.0 | 0.1 | 0.4 | 0.0 |
| ShoRAH | 3.3 | - | 3.3 | 3.3 | 4.1 | - | 2.7 | - | 3.9 | 3.4 |

Table 12: Relative edit distance of the closest reconstructed haplotype to each of the simulated HCV strains (20.000x coverage).

| | s1 | s2 | s3 | s4 | s5 | s6 | s7 | s8 | s9 | s10 | s11 | s12 | s13 | s14 | s15 |
|---|---|---|---|---|---|---|---|---|---|---|---|---|---|---|---|
| *abundance* | 2% | 2% | 2% | 4% | 4% | 4% | 6% | 6% | 6% | 8% | 8% | 8% | 13% | 13% | 13% |
| SAVAGE | 0.0 | 0.0 | 0.0 | 0.0 | 0.0 | 0.0 | 0.0 | 0.0 | 0.0 | 0.0 | 0.0 | 0.0 | 0.0 | 0.0 | 0.0 |
| Virus-VG | - | 1.1 | 0.3 | 0.2 | 0.1 | 0.0 | 0.0 | 0.1 | 0.0 | 0.0 | 0.0 | 0.0 | 0.0 | 0.0 | 0.0 |
| PredictHaplo | - | - | 0.5 | - | - | - | - | 0.5 | - | 0.2 | 0.0 | 0.0 | 0.2 | 0.0 | 0.0 |
| ShoRAH | 2.4 | 1.7 | 4.6 | - | - | - | - | - | - | - | - | - | - | 3.4 | - |

Table 13: Relative edit distance of the closest reconstructed haplotype to each of the simulated ZIKV strains (20.000x coverage).

| | strain 1 | strain 2 | strain 3 | strain 4 | strain 5 | strain 6 |
|---|---|---|---|---|---|---|
| *abundance* | 50.8% | 25.4% | 12.7% | 6.3% | 3.2% | 1.6% |
| SAVAGE | 0.0 | 0.0 | 0.0 | 0.0 | 0.0 | 0.0 |
| Virus-VG | 0.0 | 0.1 | 0.0 | 0.1 | 0.1 | 0.0 |
| PredictHaplo | - | - | - | - | - | 0.8 |

Table 14: Relative edit distance of the closest reconstructed haplotype to each of the simulated Poliovirus strains (20.000x coverage).

| | strain 1 | strain 2 | strain 3 | strain 4 | strain 5 | strain 6 | strain 7 |
|---|---|---|---|---|---|---|---|
| *abundance* | 0.5% | 1% | 2% | 5% | 10% | 20% | 61.5% |
| SAVAGE | - | - | 0.0 | 0.0 | 0.0 | 0.0 | 0.0 |
| Virus-VG | - | - | 0.0 | 0.2 | 0.0 | 0.0 | 0.1 |
| aBayesQR | - | 1.3 | 1.2 | 1.2 | 0.7 | 1.1 | 0.9 |
| PEHaplo | - | 0.8 | - | 0.8 | 0.5 | 0.3 | 0.1 |
| PredictHaplo | - | - | - | - | - | - | 0.8 |
| ShoRAH | - | - | 1.6 | 1.2 | 1.0 | 1.1 | 1.0 |

Table 15: Relative edit distance of the closest reconstructed haplotype to each of the simulated HIV pol strains at 500x coverage.

| | strain 1 | strain 2 | strain 3 | strain 4 | strain 5 | strain 6 | strain 7 |
|---|---|---|---|---|---|---|---|
| *abundance* | 0.5% | 1% | 2% | 5% | 10% | 20% | 61.5% |
| SAVAGE | - | - | 0.0 | 0.0 | 0.0 | 0.0 | 0.0 |
| Virus-VG | - | - | 0.6 | 0.0 | 0.0 | 0.1 | 0.2 |
| aBayesQR | - | 1.2 | 1.2 | 1.3 | 0.7 | 1.0 | 0.9 |
| PEHaplo | - | 1.0 | 1.1 | 0.8 | 0.3 | 0.3 | 0.2 |
| PredictHaplo | - | - | - | - | - | 0.9 | 0.8 |
| ShoRAH | - | - | - | 1.2 | 1.0 | 1.1 | 1.0 |

Table 16: Relative edit distance of the closest reconstructed haplotype to each of the simulated HIV pol strains at 1000x coverage.

| | strain 1 | strain 2 | strain 3 | strain 4 | strain 5 | strain 6 | strain 7 |
|---|---|---|---|---|---|---|---|
| *abundance* | 0.5% | 1% | 2% | 5% | 10% | 20% | 61.5% |
| SAVAGE | - | 0.0 | 0.0 | 0.0 | 0.0 | 0.0 | 0.0 |
| Virus-VG | - | 0.6 | 0.1 | 0.1 | 0.0 | 0.0 | 0.4 |
| aBayesQR | - | - | 1.3 | 1.3 | 0.9 | 1.2 | 1.0 |
| PEHaplo | 0.9 | 0.5 | 0.5 | 0.3 | 0.4 | 0.4 | 0.3 |
| PredictHaplo | - | - | - | - | - | 0.9 | 0.8 |
| ShoRAH | - | - | - | 1.2 | 1.1 | 1.3 | 1.1 |

Table 17: Relative edit distance of the closest reconstructed haplotype to each of the simulated HIV pol strains at 5000x coverage.

| | 89.6 | HXB2 | JRCSF | NL43 | YU2 |
|---|---|---|---|---|---|
| SAVAGE | 0.0 | 0.0 | 0.0 | 0.0 | 0.0 |
| Virus-VG | 0.0 | 0.0 | 0.2 | 0.0 | 0.0 |
| PredictHaplo | 1.1 | 3.3 | 1.4 | 1.5 | 1.0 |
| ShoRAH | 3.5 | 3.1 | 3.0 | 3.8 | 3.9 |

Table 18: Relative edit distance of the closest reconstructed haplotype to each of the labmix strains (excluding LTR) at 20.000x coverage.

| | 89.6 | HXB2 | JRCSF | NL43 | YU2 |
|---|---|---|---|---|---|
| **100x** | | | | | |
| SAVAGE | 0.0 | 0.0 | 0.0 | 0.0 | 0.0 |
| Virus-VG | 0.0 | 0.0 | 0.0 | 0.0 | 0.0 |
| aBayesQR | 1.6 | 1.7 | 1.5 | 1.3 | 1.8 |
| PEHaplo | 1.6 | 1.4 | 1.5 | 0.5 | 1.5 |
| PredictHaplo | - | 1.9 | - | 1.8 | 1.9 |
| ShoRAH | 1.3 | 1.4 | 1.1 | 1.2 | 1.5 |
| **1000x** | | | | | |
| SAVAGE | 0.0 | 0.0 | 0.0 | 0.0 | 0.0 |
| Virus-VG | 0.1 | 0.2 | 0.2 | 0.1 | 0.2 |
| aBayesQR | 1.7 | 1.9 | 1.4 | 1.6 | 2.0 |
| PEHaplo | 0.6 | 0.6 | 0.8 | 0.6 | 0.6 |
| PredictHaplo | 0.9 | - | 0.7 | 0.5 | 1.5 |
| ShoRAH | 1.7 | 1.6 | 1.5 | 1.3 | 1.5 |
| **20.000x** | | | | | |
| SAVAGE | 0.0 | 0.0 | 0.0 | 0.0 | 0.0 |
| Virus-VG | 0.2 | 0.0 | 0.3 | 0.1 | 0.2 |
| aBayesQR | - | - | 1.8 | 2.1 | - |
| PEHaplo | 0.2 | 0.2 | 0.3 | 0.3 | 0.5 |
| PredictHaplo | 0.4 | 0.4 | 0.4 | 0.4 | 0.5 |
| ShoRAH | 1.8 | 1.0 | 1.6 | 1.6 | 1.7 |

Table 19: Relative edit distance of the closest reconstructed haplotype to each of the labmix strains (pol-region only) at 100x, 1000x, and 20.000x coverage.

#### 6 Command lines

*Read simulations:* SimSeq
*Read trimming, adapter removal:* CutAdapt (Martin, 2011)
*Ad-hoc reference construction:* VICUNA (Yang *et al.*, 2012)
*Alignment:* BWA-MEM (Li, 2013)
*Reference-guided quasispecies reconstruction tools:* ShoRAH (Zagordi *et al.*, 2011), PredictHaplo (Prabhakaran *et al.*, 2014), aBayesQR (Ahn and Vikalo, 2018)
*De novo assembly tools:* PEHaplo (Chen *et al.*, 2018), SAVAGE (Baaijens *et al.*, 2017)
*Assembly evaluation:* QUAST (Gurevich *et al.*, 2013)

All tools were run using default settings.

**SimSeq**

```
java -jar SimSeq-master/SimSeq.jar -l 600 -1 250 -2 250 \
     -e SimSeq-master/profiles/miseq_250bp.txt -r truth.fasta \
     -n <num_reads> -o sim_reads.sam
```

**BWA-MEM** *version 0.7.15-r1140*

```
bwa mem reference.fasta reads.fastq > reads.sam
```

**VICUNA:** *version 1.3*

```
vicuna-omp-v1.0 config.txt
```

**PredictHaplo** *version 0.4*

```
PredictHaplo-Paired config.txt
```

**ShoRAH:** *version 0.8.2*

```
python shorah.py -b paired.sorted.bam -f reference.fasta
```

**aBayesQR:**

```
aBayesQR config
```

**PEHaplo:**

```
python pehaplo.py -f1 forward.fasta -f2 reverse.fasta -l 180 -r 250 -F 600 -t 8
```

**SAVAGE:** *version 0.4.0 (Bioconda)*

```
savage -p1 forward.fastq -p2 reverse.fastq --revcomp --split 30
```

**Virus-VG:**

```
python build_graph_msga.py -f forward.fastq -r reverse.fastq \
   -c contigs_stage_c.fasta -t 8 -vg vg-v1.7.0
python optimize_strains.py -m 100 -c 200 node_abundance.txt contig_graph.final.gfa
```

**QUAST:** *version 4.3*

```
python quast.py -m 500 -R ground_truth.fasta contigs.fasta
```

#### 7 Installation

Virus-VG is publicly available at `https://bitbucket.org/jbaaijens/virus-vg` under the MIT license. The repository contains detailed instructions regarding installation and dependencies. We recommend using SAVAGE for de novo assembly prior to using Virus-VG and using the Conda package manager for installing SAVAGE as well as all Virus-VG dependencies. The manual for running SAVAGE is available at `https://bitbucket.org/jbaaijens/savage`. The only dependency not (yet) available through Conda is the vg-toolkit, which has to be downloaded from `https://github.com/vgteam/vg`. Virus-VG makes use of the Gurobi Optimization software, `www.gurobi.com`, for which an academic license can be requested for free.
